# A benchmarked, high-efficiency prime editing platform for multiplexed dropout screening

**DOI:** 10.1101/2024.03.25.585978

**Authors:** Ann Cirincione, Danny Simpson, Purnima Ravisankar, Sabrina C. Solley, Jun Yan, Mona Singh, Britt Adamson

## Abstract

Prime editing installs precise edits into the genome with minimal unwanted byproducts, but low and variable editing efficiencies have complicated application of the approach to high-throughput functional genomics. Leveraging several recent advances, we assembled a prime editing platform capable of high-efficiency substitution editing across a set of engineered prime editing guide RNAs (epegRNAs) and corresponding target sequences (80% median intended editing). Then, using a custom library of 240,000 epegRNAs targeting >17,000 codons with 175 different substitution types, we benchmarked our platform for functional interrogation of small substitution variants (1-3 nucleotides) targeted to essential genes. Resulting data identified negative growth phenotypes for nonsense mutations targeted to ∼8,000 codons, and comparing those phenotypes to results from controls demonstrated high specificity. We also observed phenotypes for synonymous mutations that disrupted splice site motifs at 3′ exon boundaries. Altogether, we establish and benchmark a high-throughput prime editing approach for functional characterization of genetic variants with simple readouts from multiplexed experiments.

## Introduction

Large-scale sequencing efforts have cataloged millions of human genetic variants, including hundreds of thousands linked to human traits or diseases.^1–4^ A central challenge now is to characterize the functional effects of such variants on molecular, cellular, and physiological processes (*e.g.*, protein function, gene regulation). Technologies for multiplexed variant screening have greatly enabled such work,^5–16^ but existing approaches have limitations. For example, although ectopic gene expression can be applied in high-throughput to evaluate all possible variants across small, defined sequences,^8–11^ exogenously expressed sequences do not retain genomic context and therefore variants evaluated on such platforms do not always phenocopy their endogenous counterparts. To overcome this limitation, an approach for saturation genome editing that uses homology-directed repair (HDR) to install variant libraries into the genome at Cas9-induced DNA double-strand breaks was developed.^12,13^ This approach allows nearly any sequence change to be introduced at endogenous loci; however, variant installation with HDR can be inefficient, imprecise, and difficult to multiplex across targets,^17^ often restricting use to individual genomic regions. To further improve variant screening, base editing platforms were developed.^5–7,16^ These platforms enable efficient variant installation across the genome, but can introduce undesired bystander mutations alongside programmed edits and are restricted by mutation type (*i.e.*, cytosine base editors produce C>T or G>A edits),^14,15,18–21^ thus limiting variant scope in any individual experiment.^22–24^

An ideal platform for high-throughput variant characterization would allow precise, efficient, and multiplexable genome editing of any variant type across the genome. Approaching this ideal, prime editing can flexibly install all twelve single nucleotide substitutions, small insertions, and deletions into targeted genomic loci with minimal unintended editing.^25^ However, despite some recent high-throughput applications,^26–31^ generalized use of prime editing for multiplexed variant analysis has been limited by typically low and variable editing efficiencies. Here we show that, when applied in the absence of DNA mismatch repair (MMR) and with stably expressed and optimized editing components, prime editing is capable of efficient and precise variant installation. We then rigorously benchmark these conditions for variant screening by evaluating tens of thousands of genetic variants with expected phenotypes, demonstrating robust, high-specificity dropout effects.

## Results

### Designing a prime editing platform capable of high-efficiency editing

We sought to build and evaluate a robust prime editing platform for variant screens. The simplest form of prime editing (PE2) is a two-component system that uses an engineered Cas9 protein (Cas9 H840A nickase fused to a reverse transcriptase) and a prime editing guide RNA (pegRNA) that specifies both the DNA target and intended edit. Together, these components bind the targeted genomic locus, nick the complementary DNA strand, and reverse transcribe the intended edit into the genome at that site with few unwanted or bystander edits. Beyond the precision and flexibility of prime editing, other technical features make the approach theoretically well-suited for large, multiplexed experiments. Specifically, because all of the information required for variant installation is physically encoded in the pegRNA (*i.e.,* target and edit), the system should be compatible with standard screening protocols used for other CRISPR-based perturbation systems (*e.g.*, Cas9,^32,33^ CRISPRi/a,^34,35^ CRISPRoff,^36^ and base editors),^6,7^ including parallel synthesis of large pegRNA libraries, pooled delivery, and phenotyping by determining pegRNA frequencies across selected populations. We set out to rigorously evaluate prime editing applied in this manner.

To begin, we obtained two K562 clonal cell lines constitutively expressing different prime editor fusion proteins (PE2^25^ or PEmax)^37^ from the *AAVS1* safe-harbor locus, with EGFP co-expressed from the same transcript to enable long-term monitoring of transgene expression (herein, cell lines called PE2 and PEmax,^38^ respectively; Figures 1A and 1B). Additionally, because MMR has been shown to inhibit small prime edits,^37,39^ we generated an MMR-deficient *MLH1*-knockout derivative cell line from the PEmax cells, which we call PEmaxKO (Figures 1A and S1A). We then tested prime editing in all three cell lines with the PE2 approach at two endogenous sites (HEK3 +1 T>A and DNMT1 +6 G>C, where +1 and +6 represent the nucleotide position downstream from the Cas9 H840A nicking site) using two types of pegRNAs, those with and without a structural motif on the 3′ end (Figures 1C, 1D, and S1B). Addition of this motif, called *tevopreQ_1_*, has been shown to increase prime editing efficiencies and constitutes the standard engineered pegRNA (epegRNA) design.^40^

**Figure 1.**
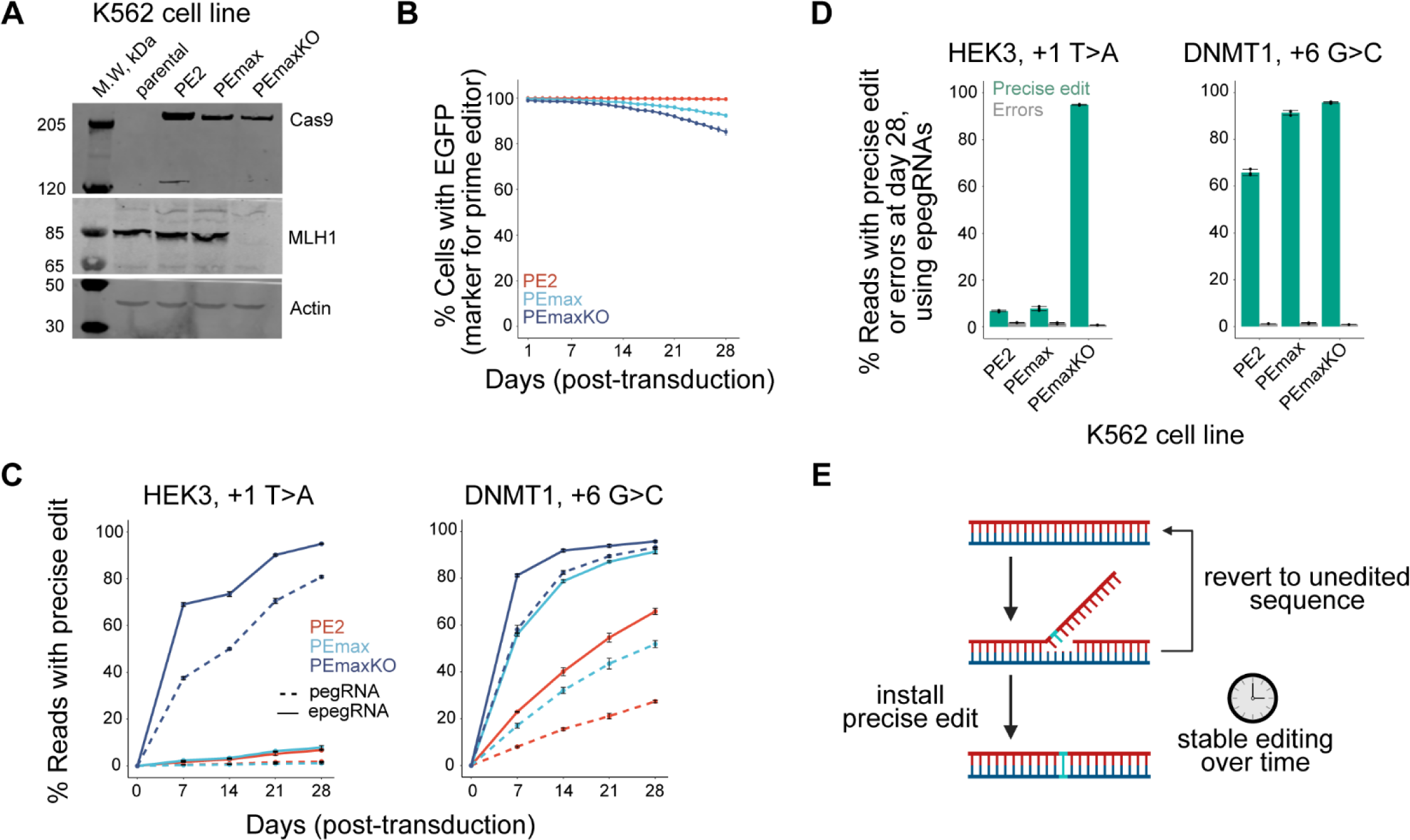
Continuous prime editing in MMR-deficient cells at two endogenous loci produces near complete installation of intended edit. (A) Western blot analysis of K562 cells (parental) and clonal derivatives stably expressing indicated prime editor protein with (PEmaxKO) or without (PE2, PEmax) genetic disruption of *MLH1*. Analysis after one month of culture post-transduction with (e)pegRNA constructs (from same cell populations as in B-D). (B) Percentages of cells with expression of marker protein (EGFP) co-expressed with prime editor protein (driven by IRES2 from the same transcript). Analysis over one month of culture post-transduction with (e)pegRNA constructs. (C) Percentages of sequencing reads containing the precise HEK3 +1 T>A (left) and DNMT1 +6 G>C (right) substitutions from cells edited with the indicated components over one month. Edits are specified using relative distance from the predicated site of the Cas9(H840A) induced nick, such that +1 indicates an edit position directly adjacent to the nick and +6 indicates an edit position 6 nt away within the protospacer adjacent motif (PAM). Day 0 represents the unedited time point, at which cells were transduced with (e)pegRNA constructs. (D) Percentages of sequencing reads containing the precise HEK3 +1 T>A (left) and DNMT1 +6 G>C (right) substitutions or errors from cells sampled 28 days after transduction of epegRNA constructs. (E) Schematic of prime editing over time, with intended edit shown in cyan. Data and error bars in B-D represent mean +/- s.d. (n=3 independent biological replicates).

Because prime editing with the PE2 approach produces primarily the intended edit or unedited sequence at the targeted site, neither of which represents an unwanted “endpoint”, stable expression of editing components should result in accumulation of precise edits over time (Figure 1E).^17,27–31,41,42^ Results from our experiments with the HEK3 +1 T>A and DNMT1 +6 G>C edits confirmed this expectation, demonstrating continuous accumulation of intended edits over one month (Figure 1C) with minimal observation of unwanted byproducts or “errors” at either site (Figures 1D and S1B). Efficiencies of both substitutions were highest in cells with the optimized prime editor (PEmax) and when using an epegRNA in the absence of MMR (Figure 1C). Indeed, this combination of features produced a remarkable ∼95% precise editing (*i.e.*, intended edit with no errors) at both sites after one month of continuous editing (Figure 1D). These results represent strong improvement over our previously reported editing frequencies for the same edits measured with transient editor expression, which did not reach higher than 30%, despite being evaluated in an MMR-deficient cell line.^37^ Directly comparing our results from PEmax and PEmaxKO cells also confirmed the benefit of MMR loss for prime editing in the context of stable editor expression. Specifically, installation of HEK3 +1 T>A with epegRNAs reached only 7.8% precise editing by day 28 in PEmax cells but reached 94.9% in MMR-defective PEmaxKO cells (Figure 1D). By contrast, installation of DNMT1 +6 G>C with epegRNAs reached high precise editing in both cell lines as early as day 14 (91.8% in PEmaxKO, 78.7% in PEmax; Figure 1C). Results with this latter edit are consistent with the observations that C-C mismatches, which are expected intermediates of G>C prime editing, are poor MMR substrates and that G>C substitutions can be efficient prime edits in the presence of MMR.^37,43–46^

### Prime editing with stable expression of PEmax and a self-targeting epegRNA “sensor” library

We next evaluated prime editing in PEmax and PEmaxKO cell lines across hundreds of edits using a self-targeting library design that links epegRNA expression cassettes to targetable “sensor” sequences (Figure 2A). Such libraries have been used previously to study prime editing and other genome editing tools^14,16,29,47–52^ and enable editing efficiencies to be quantified across many guide RNA-target pairs. To select epegRNA-target pairs for our library, we mined data from a previously published, self-targeting prime editing screen that evaluated protospacer adjacent motif (PAM)-disrupting +5 G>C edits at 2,000 target sites using 48,000 pegRNAs, including 24 pegRNA designs for each edit with different reverse transcriptase template (RTT) and primer binding site (PBS) lengths.^50^ From the 2,000 evaluated target sites in that study, we randomly selected 640. We then identified the most efficient pegRNA for each of those targets (ranging from 0.14-60.4% precise editing after 5 days) and redesigned each as three epegRNAs with identical PBS sequences and nearly identical RTTs specifying a +5 G>A, G>T, or G>C edit. Our final self-targeting library consisted of 2,000 epegRNA-target pairs (Figure S2A; Table S1), including 22 positive controls (edits tested previously at endogenous targets)^25,37^ and 58 negative controls (epegRNAs specifying the reference sequence or non-targeting epegRNAs with a scrambled target site sequence).

**Figure 2.**
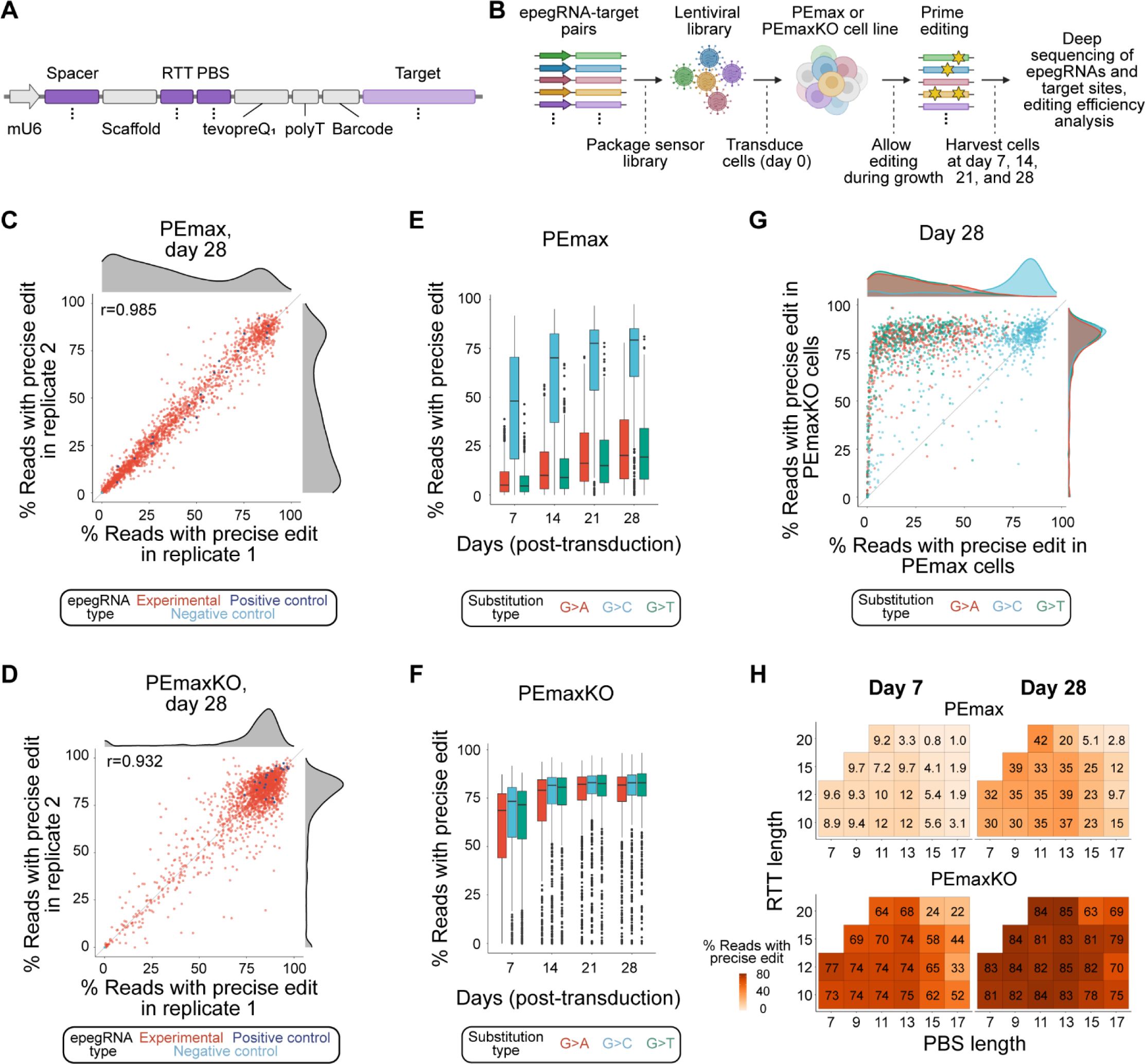
Prime editing with stable expression of PEmax and a self-targeting epegRNA “sensor” library reveals high-efficiency, precision editing. (A) Schematic of self-targeting expression cassette for sensor screens. Regions indicated with purple varied coordinately across the library (as denoted by dots), with dark purple specifying variable epegRNA components and light purple specifying the corresponding target site. mU6, modified mouse U6 promoter; RTT, reverse transcriptase template; PBS, primer binding site. (B) Schematic of workflow for sensor screens. Briefly, epegRNA-target pairs were transduced into K562 cells stably expressing PEmax with or without genetic disruption of *MLH1* (PEmaxKO or PEmax cells, respectively). Cell populations were grown for 28 days and sampled intermittently to evaluate prime editing in the target region. (C) Percentages of sequencing reads from sensor targets containing the precise edit from two replicates of a screen performed in PEmax cells and collected on day 28. Each data point represents an individual epegRNA-target pair. Correlation between replicates (Pearson’s r) indicated. Density plots on top and side show data distribution for replicate 1 and 2, respectively. (D) As in C, but for two replicates of a sensor screen performed in PEmaxKO cells. (E) Replicate-averaged percentages of sequencing reads from sensor targets containing the precise edit for experimental (non-control) epegRNA-target pairs (1,898 pairs represented in each boxplot) from a screen performed in PEmax cells and collected on indicated days. Median and interquartile range (IQR) of the full set of experimental epegRNA-target pairs installing indicated substitution types are shown. Whiskers extend 1.5*IQR past the upper and lower quartiles. (F) As in E, but for a screen performed in PEmaxKO cells. (G) Replicate-averaged percentages of sequencing reads from sensor targets containing the precise edit for experimental epegRNA-target pairs from screens performed in PEmax and PEmaxKO cells and collected on day 28. Density plots on top and side show data distribution per substitution type for PEmax and PEmaxKO cells, respectively. (H) Heatmap depicting median, replicated-averaged percentages of sequencing reads from sensor targets containing the precise edit for different RTT and PBS lengths of experimental epegRNAs (percentages listed). Data from cells collected on indicated days from indicated screens, shown for RTT/PBS combinations that were used to target at least five sensor targets.

After cloning our self-targeting library into a lentiviral vector, we transduced the epegRNA-target pairs into PEmax and PEmaxKO cells at low multiplicity of infection (MOI = 0.7), ensuring that the majority of cells received at most one pair. We then selected the transduced cells for cassette integration and grew the resulting population for approximately one month, sampling cells at 7, 14, 21, and 28 days post-transduction (Figure 2B). At each timepoint, we sequenced the epegRNA-target pairs, and after removing pairs with low read counts and unifying the data across samples, we determined editing outcomes for the remaining 1,974 pairs using a custom analysis pipeline (Figure S2B; Table S1; Methods). For each pair, we quantified three outcome categories: outcomes containing only the intended edit (“precise edits”), those with at least one error, and unedited sequence.

For many epegRNA-target pairs, we observed high-efficiency precise editing, with 20.2% (388) and 75.5% (1,453) of edits reaching 75% or higher precise editing by day 28 in PEmax and PEmaxKO cells, respectively (Figures 2C and 2D). We also observed low rates of error in all samples (median errors <4% for both cell lines on day 28; Figures S2C and S2D), and indicating strong reproducibility, precise edits and errors were well correlated across replicates for both cell lines at each time point (Pearson’s r = 0.932-0.999 for precise edits, r = 0.663-0.975 for errors; Figures 2C, 2D, S2C, and S2D). Separating results by substitution type then revealed that G>C edits outperformed the others in MMR-proficient PEmax cells (median precise editing of 79.2% for G>C, 20.2% for G>A, and 19.4% for G>T at day 28 for experimental epegRNAs) and were the majority of high-efficiency edits made in those cells (Figure 2E). Additionally, by day 14, installation of many +5 G>C edits was already high in PEmax cells, while the other substitution types were made more slowly (Figure 2E), consistent with our previous results with DNMT1 +6 G>C and HEK3 +1 T>A (Figure 1C). By contrast, each of the three substitution types were installed more synchronously on average and to high efficiencies in PEmaxKO cells (median precise editing of 83.0% for G>C, 81.8% for G>A, and 83.0% for G>T at day 28 for experimental epegRNAs; Figure 2F). Notably, +5 G>C edits performed similarly well in both cell lines by day 28 (Figure 2G). Resulting data also confirmed that, similar to previous observations,^50,52^ RTTs of 10-15 nt and PBSs of 9-13 nt generally had high rates of editing among tested epegRNAs, with or without considering G>C edits (Figures 2H and S2E).

Because the +5 G>C edits in our library were evaluated previously,^50^ we could compare editing efficiencies from our screen to those obtained in a different study with alternative conditions (transient expression of PE2 in MMR-deficient HEK293T cells using pegRNAs). For the vast majority of these edits in our self-targeting library (93.7% of edits for PEmax cells and 97.6% for PemaxKO cells; Figure S2F), we achieved higher rates of precise editing by day 28, with day 7 median efficiencies in PEmaxKO cells more than triple those previously reported at day 5 (day 7 median editing = 73.4% vs. day 5 = 20.8%).^50^ Altogether, results from our sensor screens establish potential for rapid, high-efficiency editing with stable expression of PEmax and epegRNAs using different types of PAM-disrupting edits in the absence of MMR, and for G>C edits without MMR disruption.

### Phenotype-based negative selection screening with prime editing at massive scale

We next evaluated the potential of using optimized prime editing conditions for high-throughput variant screening by evaluating a 240,000 epegRNA library specifying tens of thousands of edits intended to generate premature nonsense codons in essential genes. To design this library, we built a bespoke pipeline that identifies candidate codons, verifies that targeted bases are accessible to prime editing, filters for codons that can also accommodate synonymous mutations as controls, and then generates the epegRNA extension sequences (Figures 3A and S3A). For each of the >44,000 edits specified in this library, we included epegRNAs with up to eight different extension designs: PBS lengths of 11 nt or 13 nt and RTT lengths of 10 nt, 12 nt, 15 nt, or 20 nt. Edits were constrained to 1-3 nt substitutions positioned within 20 nts of the Cas9(H840A) nicking site (+ direction) for each protospacer. The resulting library contained 130,276 epegRNAs specifying edits designed to install nonsense codons (“stop” epegRNAs); 94,724 spacer/codon-matched epegRNAs specifying edits that do not change an amino acid and thus were not designed to alter protein function (“synonymous” controls); 12,000 epegRNAs with extension sequences specifying the reference sequence (“no edit” controls); and 3,000 epegRNAs that use non-targeting spacers (“non-targeting” controls; Figure 3B; Tables S2 and S3). The library, called StopPR (<u>stop</u> codon <u>pr</u>ime editing), targeted 17,061 codons across 1,232 commonly essential genes (defined by DepMap)^53^ and specified stop codon installation through 46 combinations of edit positions and 175 substitution types (Figure 3C).

**Figure 3.**
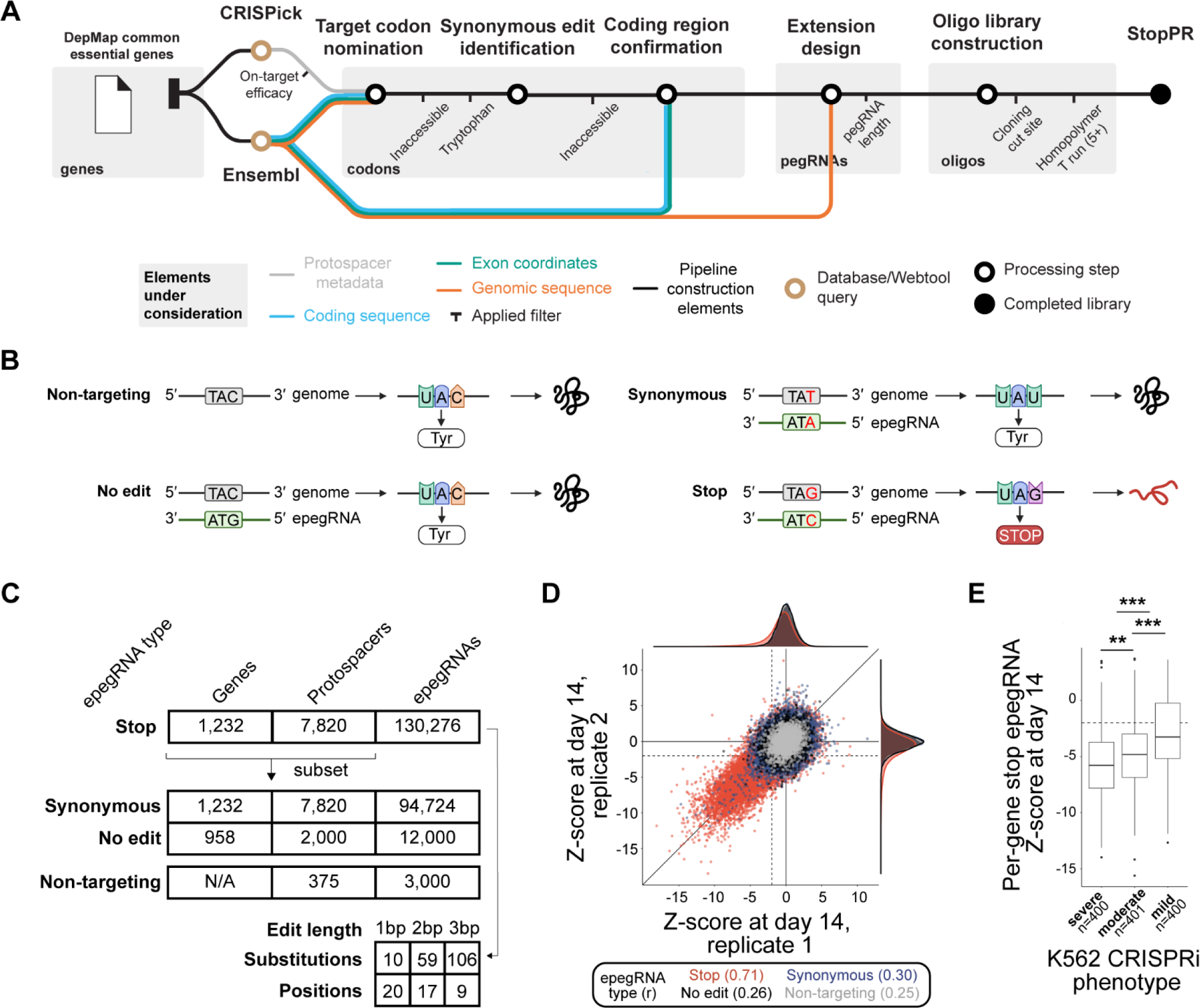
Multiplexed prime editing screen targeting premature stop codons to essential genes induces negative growth phenotypes. (A) Schematic of design pipeline used to generate StopPR epegRNA library. Pipeline used information from CRISPick^61^ and gene annotations to identify edits capable of introducing premature stop codons in essential genes. (B) Schematic illustrating the intended consequences of prime editing for each type of epegRNA included in the StopPR library: non-targeting, no edit, and synonymous controls, and stop epegRNAs. (C) Composition of StopPR epegRNA library, including numbers of each type of epegRNA and numbers of genes/protospacers targeted, as well as numbers of stop epegRNAs with different edit lengths, positions, and substitution types. Notably, multiple codons were often targeted near the same protospacer, such that 17,061 total codons were targeted with 7,820 protospacers. (D) Growth phenotypes for epegRNAs from independent biological replicates of StopPR screen collected 14 days post-transduction. Dotted lines denote phenotype cutoffs (Z < −2). Correlation (Pearson’s r) between replicates indicated for each epegRNA type. (E) Gene-level growth phenotypes from StopPR screen (calculated as the average phenotype of the absolute strongest two stop epegRNAs per gene on day 14) binned by CRISPRi phenotypes (as previously determined in K562 cells).^54^ Individual p-values were 1.13E-3 (severe vs. moderate), 4.00E-12 (moderate vs. mild), and < 2.62E-14 (severe vs. mild) from ANOVA and Tukey post-hoc (** p < 0.01, *** p < 0.001). Median and interquartile range (IQR) of the full set of epegRNAs used in this analysis are indicated. Whiskers extend 1.5*IQR past the upper and lower quartiles. Dotted line denotes phenotype cutoff (Z < −2).

To maximize editing efficiency across substitution types, we screened our StopPR library in our PEmaxKO cell line (Figure S3B). Briefly, after transducing PEmaxKO cells (MOI = 0.7) with the library, we selected them for cassette integration and grew the resulting population for approximately one month, sampling cells at 7, 14, and 28 days post-transduction. To determine growth phenotypes, we sequenced the integrated epegRNAs from each sample (Figure S3C), removed epegRNAs with low read counts as well as a small number of epegRNAs that failed an updated design filter (including a few targeting introns; Methods), and calculated log_2_ fold changes in relative abundance at days 14 and 28 compared to day 7, which we then expressed as Z-scores normalized to the distribution of non-targeting control phenotypes (Tables S2 and S3; Methods). Altogether, we recovered growth phenotypes for 106,092 pairs of stop and synonymous epegRNAs (covering 91.7% of targeted codons), 10,007 no edit controls, and 2,312 non-targeting controls.

Using Z < −2 to threshold growth phenotypes, we found that 17.1% (18,187) of stop epegRNAs induced a negative growth phenotype by day 14 (Figure 3D), increasing to 23.1% (24,510) by day 28 (Figure S3D), but relatively few phenotypes were observed among controls (2.3%, 2,024 epegRNAs across all sets of controls by day 28). Additionally, phenotypes for stop epegRNAs were correlated between replicates (Pearson’s r = 0.71 at day 14), indicating that the observed phenotypes were reproducible, while measurements across all control epegRNAs showed little correlation between replicates, as expected (Pearson’s r = 0.29 at day 14; Figures 3D and S3D). Notably, among stop epegRNAs, our rates of negative phenotype induction were encouragingly similar to results from a base editing screen that reported 30.5% of sgRNAs designed to install nonsense codons in essential genes caused negative growth phenotypes.^7^

Within our StopPR library, individual codons were targeted by multiple stop epegRNAs with different design features (average of ∼7 epegRNAs per codon analyzed for growth phenotypes). Examining the relative potency of stop epegRNAs targeting the same codons revealed that resulting phenotypes often varied in strength (Table S3), presumably due to differences in epegRNA activity. However, of codons analyzed for growth phenotypes, 40.6% (6,353 of 15,646) were associated with a negative phenotype (Z < −2) from one or more stop epegRNAs by day 14, a rate that increased to 50.8% (7,948) by day 28. These results demonstrate that over half of our stop epegRNAs would have produced the expected phenotype had the library included only the most active designs per targeted codon. We next used our data to generate gene-level phenotypes (Methods) and found that the StopPR screen successfully reported 80.1% (984 of 1,228) of targeted genes as required for cell growth by day 14, improving to 89.3% (1,097) by day 28. Comparing these phenotypes to results from a published CRISPRi screen^54^ (also performed in K562 cells) showed general agreement with phenotypic strength (Figure 3E). Altogether these results establish that prime editing can perturb genes in high-throughput with enough efficiency to generate reproducible dropout phenotypes without sequencing the edited locus.

Because the majority of phenotypes induced by our stop epegRNAs were observed by day 14, we concluded that editing efficiencies for those targets were sufficient to impact cell fitness by that time, which is consistent with high levels of editing observed in our self-targeting screen in PEmaxKO cells at 14 days (Figure 2F). However, while Z-scored negative growth phenotypes of stop epegRNAs were correlated across time points (Pearson’s r = 0.73; Figure S3E), epegRNAs with the strongest phenotypes at day 14 (Z < −10; 244 epegRNAs) showed weaker phenotypes on day 28. This effect reflects increased noise in non-targeting controls after longer experimental time without a correspondingly strong decrease in stop epegRNA abundances at the later timepoint, which for some epegRNAs could be driven by near total population depletion by the earlier time point. Taken together, these data offer insight into the impact of experimental length and suggest that two weeks is sufficient to carry out phenotype-based screens in this context.

### Features of epegRNA design and targeted loci that influence phenotype

We next asked how aspects of epegRNA design (*e.g.*, PBS and RTT length, edit position) and characteristics of the targeted genomic loci (*e.g.,* edit location within gene bodies) impacted phenotype. For each feature, we first defined relevant groups of stop epegRNAs (*e.g.,* those targeting the beginning, middle, or end of a gene). Within these feature groups, we included only the top two epegRNAs per gene with the absolute strongest growth phenotypes, excluding epegRNAs that disrupted the PAM (*i.e.*, introduced edits at position +5-6) from all groups except edit position type. These subsettings were intended to enrich for functional epegRNAs while also ensuring equal representation of epegRNAs targeting all genes in our library and avoiding expected strong effects from edit position on other features. We then compared the average growth phenotype between combinations of those feature groups by calculating the Cohen’s d effect size. This effect size measurement describes the difference between the means of two groups normalized to the standard deviation of the underlying data, with those greater than 0.8 in magnitude generally being considered “large”.^55^ Results showed that many features had a mild effect on phenotype (Figures 4A and S4A), with the strongest effect from edit position relative to the Cas9(H840A) nicking site. Consistent with previous reports,^50–52^ edits installed in the invariant GG nucleotides of the PAM sequence for Cas9 (positions +5-6) generally resulted in stronger phenotypes than other positions, with prior positions (+1-4) remaining effective to a lesser degree, and further positions (+7-20) typically less effective, particularly beyond the +16 position (Figure 4B). While +7-20 edits were overall less effective, such edits still comprised 24.9% of stop epegRNAs with Z < −2 at day 14 (compared to being 39.1% of all analyzed stop epegRNAs). We also found that edit location within the targeted gene body and orientation of epegRNA spacer sequence with respect to gene expression contributed to phenotypic strength (Figures 4A and S4A). Specifically, we observed that edits toward the beginning of genes were more disruptive, which could be attributed to higher editing efficiency earlier in the gene (as has been previously reported),^42^ but also may reflect contribution from nonsense-mediated mRNA decay before the last exon of a gene.^56^ Similarly, epegRNAs with antisense spacers (Figure S4B) were slightly more effective (Figure S4C). This observation could be due to higher editing efficiency on the template strand or a higher phenotypic impact of those edits from earlier disruption of functional protein expression.

**Figure 4.**
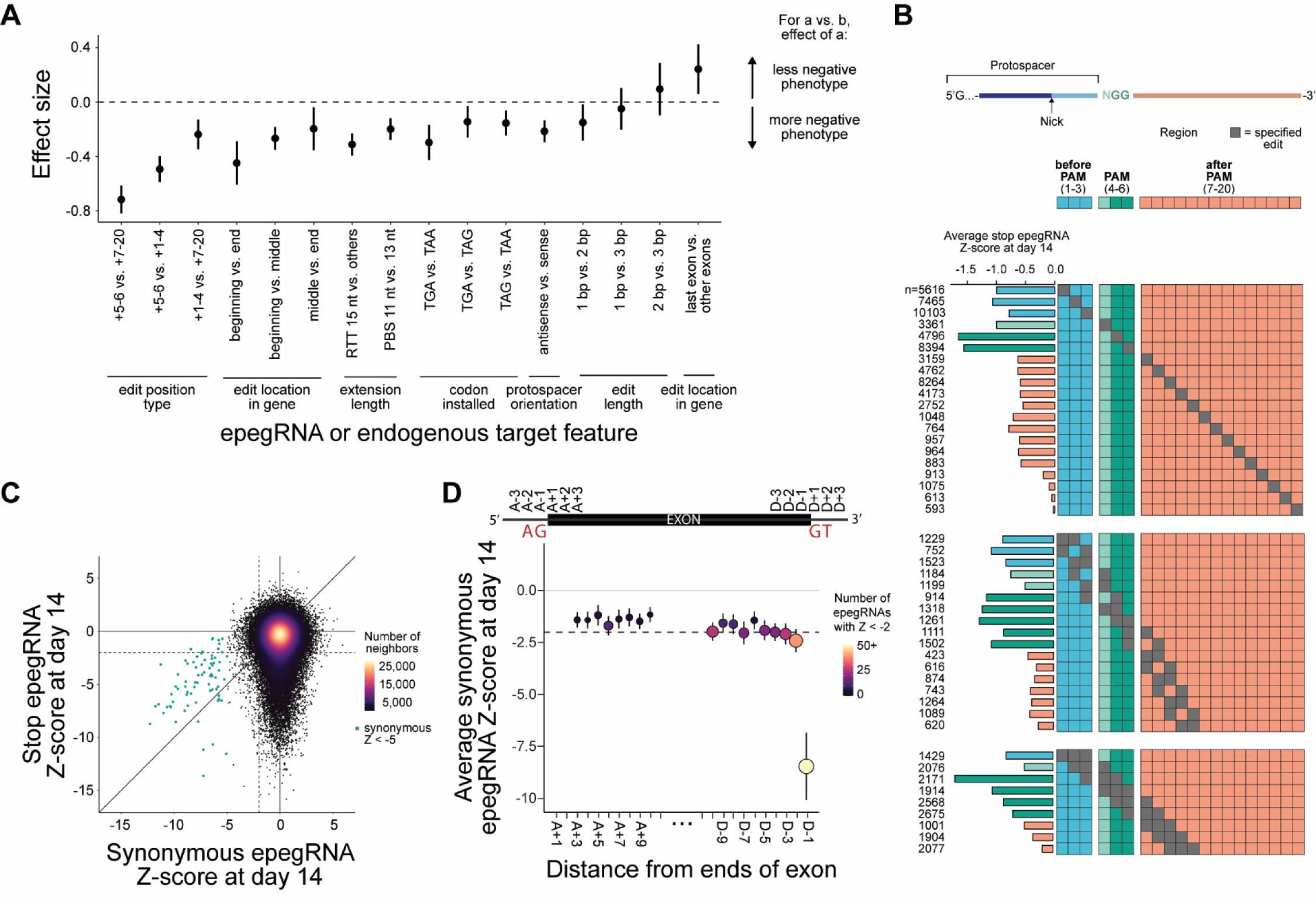
Identification of epegRNA and endogenous target features that influence prime editing-induced phenotypes. (A) Cohen’s d effect size for various aspects of epegRNA design and characteristics of targeted genomic loci. All features except edit position type were evaluated without +5-6 edits. Top two stop epegRNAs with the absolute largest phenotypes at day 14 analyzed per gene. For edit location in gene, beginning, middle, and end refer to editing within (0-33]%, (33-67]%, or (67-100]% of a gene, respectively. For extension length, other RTT lengths include 10 nt, 12 nt, and 20 nt. Bar ranges indicate 95% confidence intervals, so that intervals including effect size 0 are not significant. All p-values are listed in Figure S4A. (B) Average growth phenotypes from StopPR screen sampled from day 14 for stop epegRNAs with edits specified in the same positions (dark gray). Colors designate four position ranges. Blue indicates positions +1-3 with respect to the single-stranded nick (before PAM), light green indicates +4 (PAM-N), dark green indicates +5-6 (PAM-GG), and peach indicates +7-20 (after PAM). Numbers of stop epegRNAs denoted (left). (C) Replicate-averaged growth phenotypes for stop and spacer/codon-matched synonymous epegRNAs from StopPR screen sampled from day 14. Data points colored by density, indicated by number of neighbors. Dotted lines denote phenotype cutoffs (Z < −2). Green dots indicate strong negative growth phenotypes (Z < −5) associated with 69 synonymous epegRNAs. (D) Growth phenotypes for synonymous epegRNAs (bottom plot) from StopPR screen sampled from day 14, binned by position relative to exon boundaries (top schematic). Phenotypes were calculated as the average of 50 epegRNAs with strongest negative phenotype at each position. Positions A+1 and A+2 were excluded, as fewer than 50 synonymous epegRNAs targeted those positions. Vertical lines indicate 95% confidence intervals generated for each average. Horizontal dotted line denotes phenotype cutoff (Z < −2). Splice site acceptor (AG) and donor (GT) motifs indicated in schematic.

While informative, evaluation of epegRNAs by feature grouping is potentially confounded by the fact that epegRNAs within any such group unavoidably represent multiple features. For example, if not equally distributed across relevant categories, epegRNAs specifying edits at the beginning of genes could influence the average effect size among epegRNAs with antisense spacers. To evaluate the effect of many features simultaneously (*i.e.*, those enumerated in Figure 4A plus substitution type and targeted codon), we built a multiple linear regression model with scaled input features so that the resulting beta coefficients rank the relative impact of each feature on phenotype (Methods). Results from this model confirmed that each of the features previously interrogated by effect size contributed to growth phenotypes, with RTT length, edit position relative to the Cas9(H840A) nicking site, substitution type, and edit location within the gene body having the strongest effects (Table S4).

Subsetting our screen results by the most important features found by our model (*i.e.*, RTT of 15 nt, editing positions +5 and/or +6, and targeting codons within the first 33% of genes with the top 25% of substitution types determined to have the strongest impact on phenotype) more than doubled our day 14 rate of phenotype induction (39.3% of 1,969 stop epegRNAs; Z < −2) compared to the overall library (17.1%). Expanding this subset to include epegRNAs targeting positions +1-6 (4,021 epegRNAs total) also increased the rate of phenotype induction (by 1.9x to 32.7%). These results confirm the importance of (e)pegRNA design for phenotype-based prime editing screens and identify features and considerations that can be used in the design of future screens.

We next investigated the potential effects of chromatin context on phenotype induction using the recently released ePRIDICT tool.^57^ We determined ePRIDICT scores for nearly all codons targeted in our StopPR library (Methods), and used published score thresholds to identify those with favorable (“high” ePRIDICT score, > 50) or unfavorable (“low” ePRIDICT score, < 35) chromatin contexts. Of the 15,008 codons targeted by StopPR with ePRIDICT scores, 35.8% (5,378) were classified as favorable while just 0.9% (138) were unfavorable. Moreover, 74.0% (11,106) were in the highest 25% of all ePRIDICT scores genome-wide, indicating a better-than-average chromatin context for prime editing. This uneven distribution of scores likely reflects the pan-essentiality of genes targeted by our library, which we expect to be expressed and thus positioned in favorable chromatin contexts. Nevertheless, we observed an enrichment for phenotype induction among stop epegRNAs targeting codons with favorable scores (Odds Ratio (95% confidence interval) = 1.87 (1.81-1.94); Fisher’s exact test p-value = 2.36E-301) and mild depletion among those with unfavorable scores (OR (95% CI) = 0.81 (0.67-0.97); Fisher’s exact test p-value = 0.02), demonstrating that ePRIDICT has potential to aid epegRNA library design even when targeting generally favorable regions of the genome. Indeed, restricting our StopPR library to only targets with high ePRIDICT scores showed a 35% increase in our phenotype induction rate at day 14 (23.1%; 8,373 of 36,223 stop epegRNAs).

### Phenotypes induced by prime editing are highly specific

For each of the 130,276 stop epegRNAs within our StopPR library, we designed and screened one spacer/codon-matched synonymous control. These controls specified synonymous edits at the same codon as their matched stop epegRNAs. Similar to no edit and non-targeting controls, this subset of epegRNAs demonstrated very low activity (2.4% with Z < −2 at day 14), and associated results showed low correlations between replicates (Pearson’s r = 0.30 at day 14; Figure 3D). Critically, even at codons where nonsense mutations produced strong phenotypes (4,090 stop epegRNAs with Z < −5), we observed few effects from synonymous epegRNAs (119 with Z < −2; Figure 4C). This low incidence of unintended phenotypes indicates extremely high specificity for growth phenotypes attributed to stop epegRNAs. Moreover, this rate of unintended phenotypes compares favorably to other platforms developed for variant screening. For example, using the same cutoff (Z < −2), base editing screens previously identified 8.7-26.5% of sgRNAs designed to install silent edits with no phenotypic effect in essential genes as detrimental to growth.^7^ Similarly, a platform built with the more indel-prone PE3 approach to prime editing showed that 7.9-11.2% of no edit controls significantly depleted in growth screens.^27^ These comparisons demonstrate that, while false positives can be relatively more common on other platforms, presumably due to unintended on-target edits or reproducible off-target effects, our approach achieves high specificity.

### Unbiased identification of splice site variants

While only a minority of synonymous epegRNAs produced growth phenotypes in our screen, among those that did, we observed a set that caused unexpectedly strong effects (69 targeting 25 loci with Z < −5), including 61 that induced a stronger phenotype than the corresponding spacer/codon-matched stop epegRNA (green dots in Figure 4C). Further investigation revealed that the vast majority of these epegRNAs (65 targeting 23 loci) disrupted splice site motifs at 3′ exon boundaries, and manual inspection of sequences at the remaining two loci revealed that one specified an edit adjacent to a potential cryptic splice donor site (Figure S4D). Given these results, we reevaluated the edit locations of all synonymous epegRNAs in our StopPR library. We found that 2,637 targeted the last nucleotide at the 3′ end of an exon (D-1 position), with slight overrepresentation of that edit position due to the presence of a PAM within the canonical splice site donor motif (<u>AGG</u>T).^58^ Among this subset of epegRNAs, we observed an enrichment for negative growth phenotypes (11% with Z < −2, compared to 1.6-3.4% for epegRNAs targeting nearby exonic positions and 2.4% for the full set of synonymous epegRNAs). Moreover, examining effects from the strongest epegRNAs at positions relative to exon boundaries (within 10 bp of either boundary; 50 epegRNAs per position) revealed no strong phenotypes (Z < −5) at nearby positions (Figure 4D). Notably, reexamining epegRNAs designed as synonymous but subsequently excluded from analysis due to an updated design filter revealed that a very small number (68) targeted the intronic base immediately adjacent to D-1 (D+1 position; Methods). Among these epegRNAs, we found a similar enrichment of negative growth phenotypes (22.1%), consistent with mutational intolerance at the D-1 and D+1 positions, which has previously been observed through analysis of naturally occurring near-splice-site mutations.^59^ These results demonstrate use of our platform for interrogation of an additional class of genetic variants.

## Discussion

The development of prime editing has sparked wide interest in its potential use for high-throughput characterization of genomic variants. Evident of that interest, several groups have recently reported or preprinted proof-of-principle prime editing screens (Table S5).^26–31^ While each of these efforts represents an informative step forward, each has relied on at least one of the following experimental features to circumvent low prime editing efficiencies: positive selection phenotypes,^28^ readouts filtered by or calculated from editing efficiencies measured with exogenous “sensors” or endogenous target sequencing,^26,29–31^ or a more efficient but indel-prone version of prime editing (PE3).^26,27^ Screening platforms demonstrated by other studies are thus limited in some capacity: positive selection screens are restricted to specific phenotypes; screens that rely on sensors increase cost and experimental complexity; platforms that calculate phenotypes from endogenous target sequencing cannot be easily multiplexed across genetic loci; and PE3 increases unintended editing at targeted sites, potentially confounding results.

We sought to develop an accurate and generalizable prime editing screening platform that could be used with standard screening protocols (*e.g.*, parallel synthesis of epegRNAs, phenotypes calculated from epegRNA abundance). To build this platform, we implemented PE2-based prime editing with stable expression of PEmax and epegRNAs in MMR-deficient cells. We found that, when implemented with these features, prime editing can install precise variants with high efficiency (across hundreds of PAM-disrupting edits) and can generate reproducible growth-based dropout phenotypes in large, pooled experiments, achieving a high rate of phenotype induction across codons in essential genes targeted with nonsense mutations (50.8% of codons by day 28 with at least one epegRNA out of multiple tested). Additionally, phenotypes from splicing variants not intentionally designed into our library revealed strong potential for discovery-based applications. Nevertheless, two features of our platform, MMR-deficiency and stable expression of editing components, may continue to impede some applications as they necessitate cell engineering prior to screening. As a potential solution to at least one of these requirements, we found that +5 G>C substitutions are typically installed with high efficiency with and without MMR, suggesting that such edits may be suitable for applications where MMR cannot be inactivated.

To benchmark our approach for negative selection screening, we evaluated a highly complex prime editing library comprising hundreds of thousands of epegRNAs, representing the largest library used for phenotype-based prime editing screening to date by an order of magnitude (Table S5). Concurrent studies piloting alternative platforms have performed similar (albeit much smaller) benchmarking experiments using variants of known function. For example, a recent study that applied the PE3 approach to variant screening examined growth phenotypes from 115 epegRNAs specifying stop codons in growth-related genes and observed reproducible effects;^27^ however, correlated phenotypes from spacer-matched, no edit controls suggest that non-specific effects may confound interpretation of results with that approach. Another effort examined dropout phenotypes from nonsense mutations targeted to one essential gene (*RPL15*) with the PE2 approach and found that variant effects were categorized as depleting more often when endogenous site sequencing was used to determine phenotypes than without (80% of nonsense variants called as detrimental while only 32% were called using epegRNA abundances; 25 total variants targeted).^30^ When evaluating the same gene in our StopPR screen, we observed negative growth phenotypes (Z < −2) for 77.8% (7/9) of targeted nonsense variants with at least one stop epegRNA without endogenous target site sequencing. Our results therefore compare favorably to contemporary platforms and demonstrate the ability to measure high-specificity and reliable phenotypes from epegRNA abundance alone.

A key challenge moving forward will be to increase the efficiencies of prime editing libraries overall, thus enabling screening with fewer epegRNAs per target. Our results highlight the importance of this goal, as 23.1% of stop epegRNAs from our StopPR library induced negative growth phenotypes by day 28. While promising and comparable to rates observed using base editor technology, this rate of phenotype induction could be improved. To aid construction of more active epegRNA libraries, we identified features of epegRNA design and targeted loci that contribute to activity (*e.g.*, edit location in the gene body), although some of these features may also limit target selection. Additionally, our growth phenotypes and the tens of thousands of epegRNAs responsible for them should provide a useful resource for efforts to develop and test new prime editing tools, including experimental systems, computational pipelines, and analytical approaches, which could further improve screening.

In sum, we demonstrate the first proof-of-principle for conducting precise, massively parallel dropout screening with prime editing using standard screening protocols and the highly specific PE2 approach. We also robustly benchmark this approach to help enable high-throughput applications of prime editing in the future.

## Acknowledgements

We thank members of the Adamson lab for feedback and support throughout the course of this project. We thank former lab member K. Tam for helpful discussions. We thank W. Wang, J. Miller, and J.A. Volmar (Genomics Core Facility of Princeton University), as well as T. DeCoste and K. Rittenbach (Flow Cytometry Core Facility of Princeton University). Figures 1E, 2A, 2B, 3B, S3A, and S3B were created with BioRender.com. Research was supported by the National Institutes of Health under award numbers R35GM138167 (B.A.), RM1HG009490 (B.A.), R01-GM076275 (M.S.), P30CA072720 (Rutgers Cancer Institute of New Jersey via an National Institutes of Health Cancer Center Support Grant), T32HG003284 (Princeton QCB training grant), and T32GM007388 (Princeton MOL training grant), as well as Princeton University (B.A.), the Searle Scholars Program (B.A.), and Princeton Catalysis Initiative (B.A.). A.C. was supported by the National Science Foundation Graduate Research Fellowship Program (DGE-2039656). J.Y. was supported by a fellowship provided by the China Scholarship Council (CSC), based on the April 2015 Memorandum of Understanding between the CSC and Princeton University.

## Author contributions

Conceptualization, B.A.; Methodology, A.C., D.S., J.Y., and B.A.; Generation of cell lines, P.R. and J.Y.; Library design, A.C. and D.S.; Library cloning, A.C., D.S., P.R., and S.C.S.; Western blot, S.C.S.; Screening, A.C. P.R., and S.C.S.; Pipeline development, D.S.; Formal analysis, A.C. and D.S.; Writing, A.C., D.S., and B.A., with input from all authors; Supervision, M.S. and B.A.; Project administration, B.A.; Funding acquisition, M.S. and B.A.

## Inclusion and diversity

One or more of the authors of this paper self-identifies as a gender minority in their field of research. One or more of the authors of this paper self-identifies as a member of the LGBTQIA+ community.

## Methods

### Resource availability

#### Lead contact

Further information and requests for resources and reagents should be directed to and will be fulfilled by the Lead Contact, Britt Adamson.

#### Material availability

Both epegRNA libraries (lDS004 and lAC002, also referred to as StopPR) generated in this study will be deposited to Addgene. Cell lines will be available upon request.

#### Data and code availability

Processed data from both screens are available as supplementary tables to this manuscript (Tables S1, S2, and S3). Raw sequencing data from all screens will be deposited to the NCBI GEO repository. Scripts used to process data from the self-targeting screen are available on Github at https://github.com/simpsondl/TSpeg. Scripts used to process data from the StopPR screen and generate manuscript figures will be available on Github at https://github.com/anncir1/StablePE.

### Experimental model and subject details

#### Prime editing cell lines

All prime editor constructs contained an SpCas9(H840A) nickase, fused to an MMLV RT (D200N, T306K, W313F, T330P, L603W). In addition, PEmax editor construct contained a codon-optimized MMLV RT and the following additional mutations in the SpCas9 nickase: R221K and N394K. Construction of PEmax cell line described previously.^38^ PE2 cell line constructed in the same manner as PEmax cell line. To construct *MLH1* knockout PEmax cells (PEmaxKO), 122 pmole Alt-R S.p. Cas9 Nuclease V3 (IDT 1081058) and 200 pmole Alt-R CRISPR-Cas9 sgRNA targeting *MLH1* (IDT Hs.Cas9.SSB.1.AA, 5′- mC*mU*mU*rCrArCrUrGrArGrUrArGrUrUrUrGrCrArUrGrUrUrUrUrArGrArGrCrUrArGrArArArUrArGrC rArArGrUrUrArArArArUrArArGrGrCrUrArGrUrCrCrGrUrUrArUrCrArArCrUrUrGrArArArArArGrUrGrGr CrArCrCrGrArGrUrCrGrGrUrGrCmU*mU*mU*rU) were complexed for 20 minutes at room temperature and were nucleofected into 5E5 PEmax parental cells using the SE Cell Line 4D-Nucleofector X Kit (Lonza V4XC-1032) and program FF-120, according to the manufacturer’s protocol. 5 days post nucleofection, cells were sorted by BD FACSAria Fusion Flow Cytometer into 96-well plates at 1 cell per well with 150 μL conditioned culture medium. Single cells were grown and expanded for 2-3 weeks into clonal lines. Clones with a high percentage of cells with expression of EGFP according to AttuneNXT flow cytometry analysis were selected for further characterization.

#### General cell culture and selection conditions

Lenti-X 293T was purchased from Takara (632180) and K562 (CCL-243) was purchased from ATCC. K562 stable prime editing cell lines were maintained in RPMI 1640 medium (Gibco) supplied with 10% FBS (Corning) and penicillin/streptomycin (Gibco, 100 U/mL). HEK293T cells were maintained in DMEM medium (Corning) supplied with 10% FBS and penicillin/streptomycin. All cells were kept in a humidified incubator at 37°C, 5% CO_2_. For pooled screens, K562 cells were kept in a humidified multitron at 37°C, 5% CO_2_, 52-76 rpm depending on total volume.

#### General sequences and cloning

For endogenously tested HEK3 +1 T>A and DNMT1 +6 G>C substitutions, spacer and 3′ extension sequences were from a previous publication (HEK3_4a_1TtoA and DNMT1_ED5f _6GtoC, respectively),^25^ modified scaffold sequence was 5′GTTTAAGAGCTATGCTGGAAACAGCATAGCAAGTTTAAATAAGGCTAGTCCGTTATCAACTTGAAAAAGTGGCACCGAGTCGGTGC,^60^ and RNA structural motif for epegRNAs was *tevopreQ_1_* (5′-CGCGGTTCTATCTAGTTACGCGTTAAACCAACTAGAA).^40^ pegRNAs and epegRNAs used the pU6-sgRNA-EF1Alpha-puro-T2A-BFP (Addgene #60955)^35^ backbone. Cloning details for these guides described previously.^38^

To create a backbone plasmid suitable for use in cloning our self-targeting epegRNA library (lDS004), an intermediate backbone plasmid (pJY126) was first generated by removing BsmBI restriction sites on pU6-sgRNA-EF1Alpha-puro-T2A-BFP (Addgene #60955)^35^ through Golden Gate Assembly (NEB E1602S). Then, through restriction cloning, a DNA duplex annealed from DNA oligos (5′-TTGGGAGACGCCTGCAGGCTGCTAAGCTAGGCGCGCCCGTCTCATTTTTTTC, 5′-TCGAGAAAAAAATGAGACGGGCGCGCCTAGCTTAGCAGCCTGCAGGCGTCTCCCAACAAG) was inserted into pJY126 digested with BstXI and XhoI. This intermediate backbone (pJY127) was then digested with BamHI (NEB R0136S) and NotI (NEB R0189S), and a DNA duplex annealed from DNA oligos (5′-GATCCAGATCGGAAGAGCACACGTCTGAACTCCAGTCACGC, 5′-GGCCGCGTGACTGGAGTTCAGACGTGTGCTCTTCCGATCTG) was inserted through restriction cloning to produce the final pAC025 backbone plasmid.

To create a backbone plasmid suitable for use in cloning our StopPR library (lAC002) by including a *tevopreQ_1_* motif,^40^ we first inserted a DNA duplex annealed from DNA oligos (5′-CGCGCCCGTCTCACGCGGTTCTATCTAGTTACGCGTTAAACCAACTAGAATTTTTTTC, 5′-TCGAGAAAAAAATTCTAGTTGGTTTAACGCGTAACTAGATAGAACCGCGTGAGACGGG) into pJY127 digested with AscI (NEB R0558S) and XhoI. This intermediate backbone (pJY128) was then digested with BamHI and NotI, and a DNA duplex annealed from DNA oligos (5′-GATCCAGATCGGAAGAGCACACGTCTGAACTCCAGTCACGC, 5′-GGCCGCGTGACTGGAGTTCAGACGTGTGCTCTTCCGATCTG) was inserted through restriction cloning to produce the final pAC026 backbone plasmid.

### Method details

#### Western blot for prime editor and MLH1

Cells were harvested from cell culture (1E4 cells µL^-1^) and lysed in 1x Lysis buffer (1x NuPage LDS, 50 mM Sample Reducing Agent). After resuspension via vortex, samples were incubated at 70°C for 10 min. Temperature was raised to 85°C for 3 min. After incubation, samples were moved to room temperature and Benzonase Mix (final concentration 5 mM MgCl2, 1.25 U µL^-1^ benzonase) was added. Samples were then incubated at 37°C for 30 min and subsequently used for protein electrophoresis. Samples (1E5 cells) were loaded and run on 3-8% Tris-Acetate Gels (ThermoFisher) in Running Buffer (1x NuPage Tris-Acetate Running Buffer, 2.5x NuPage Antioxidant) at 180 V until completion. Proteins were then transferred to an ethanol-activated PVDF membrane (BioRad) in Transfer Buffer (1x NuPage Transfer Buffer, 10% Methanol, 2.5x NuPage Antioxidant, 0.025% SDS) at 30 V for 1 hr. Protein transfer and total protein content was assessed by Ponceau Staining (Sigma Aldrich). Ponceau Stain was washed out with 1x TBST, and then membranes were incubated in Blocking Buffer (1x TBST and 5% Dry Milk) for 1 hr at room temperature. Membranes were then incubated overnight on a shaker at 4°C in primary antibodies (ß-actin CST3700S; MLH1 Invitrogen MA5-32041; Cas9 Takara 632607) diluted 1:1000 in 1x TBST with 3% BSA, washed 3x in 1X TBST for 5 min, and then incubated in secondary antibody (1x Licor Intercept Buffer, 1:20000 IRDye Secondary Antibodies) for 1 hr at room temperature in dark. Before imaging on a Li-Cor Odyssey Infared Imaging system, membranes were washed 3x in 1x TBST for 5 min.

#### Oligonucleotide library designs

##### Self-targeting library (lDS004)

640 target sites in human protein-coding genes were randomly selected from “library 1” in Kim et al.^50^ and the corresponding highest-efficiency RTT/PBS length combination was determined for each selected site. We then designed three epegRNAs per target site with the selected PBS and identical or nearly identical RTT sequence, each specifying a +5 G>A, G>T, or G>C edit. With the addition of 22 positive control epegRNAs for sites tested endogenously in the literature, 51 non-targeting controls (with a scrambled target site sequence), and 7 no edit controls (with epegRNAs specifying the reference sequence), the final library of 2,000 epegRNA-target pairs tests seven PBS lengths (7, 9, 11, 13, 14, 15, and 17 nt), nine RTT lengths (10, 11, 12, 13, 14, 15, 17, 20, and 22 nt), and all three G>N mutations at the +5 position (Table S1).

epegRNAs and accompanying target sites were synthesized as 250 nt oligonucleotides by Twist Bioscience. Oligonucleotides were structured with adaptor sequences on both ends for library amplification, specifically 5′-GTATCCCTTGGAGAACCACCT on the 5′ end and 5′-CAGACGTGTGCTCTTCCGAT on the 3′ end, with internal BstXI (5′-CCACCTTGTTGG) and BamHI (5′-GGATCC) restriction enzyme sites surrounding epegRNA components (19 nt sgRNA and 17-39 nt extension sequences, 37 nt *tevopreQ_1_*^,40^ and 7 nt poly-T), 17 nt barcodes unique to each epegRNA-target pair, and 45 nt target sites, with reversed BsmBI restriction enzyme sites (5′-GTTTAGAGACGGCATGCCGTCTCGGTGC) splitting the sgRNA target sequence from the remainder of designed components to facilitate a two-step cloning process. Target sites were designed to include 4 nt upstream of the protospacer sequence in addition to the PAM and full RTT binding site.

##### StopPR (lAC002)

A set of 1,247 genes were nominated for inclusion in StopPR due to their determined status as common essential genes by DepMap.^53^ CRISPick^61^ was used to design 35 sgRNAs targeting each gene using reference genome Human GRCh38 (NCBI Refseq) with CRISPRko and SpyoCas9 options, which were then filtered to 16,278 sgRNA target sequences with on-target efficacy scores > 0.5. Ensembl Biomart^62^ was used to obtain exon coordinates, coding sequences, and full genomic regions for each target gene. Codons accessible to each protospacer that could be mutated to stop codons with 1 bp, 2 bp, or 3 bp mutations were identified, then any edits which could not be targeted with prime editing were removed; this latter case could occur when the Cas9 cut site occurs within the targeted codon. For each targeted codon, mutations inducing a synonymous amino acid change (such as mutating the codon ACA to ACG, both encoding threonine) were also identified, and codons where the synonymous mutation could not be introduced were filtered, including the removal of all tryptophan codons, as only one codon sequence produces it. For each edit, we designed accompanying PBS (11 nt, 13 nt) and RTT (10 nt, 12 nt, 15 nt, 20 nt) sequences, and filtered any combinations which would result in a too-long oligonucleotide for synthesis.

epegRNA sequences were then designed into 120 nt oligonucleotides with flanking 5′ (5′-CACCAGAAGCCACCTTGTTG) and 3′ (5′-CTGTGTTGGTCTCCCGCG) amplification regions containing BstXI and BasI restriction enzyme sites for synthesis by Twist Bioscience. sgRNA and extension sequences were split by reversed BsmBI restriction enzyme sites (5′-GTTTAGAGACGGCATGCCGTCTCGGTGC) to enable a two-step cloning process. Finally, oligonucleotides which contained incidental restriction enzyme sites or homopolymer T runs (5+) were removed, and 12,000 epegRNAs designed to introduce no edits with additional 3,000 epegRNAs containing scrambled non-targeting spacer sequences were included to generate a library of 240,000 epegRNAs (Tables S2 and S3). Notably, during later analysis of data generated with the StopPR library, an updated design filter identified a small number of epegRNAs with erroneous features (580 pairs of spacer/codon-matched stop and synonymous epegRNAs for which either epegRNA was affected). These were removed prior to all analysis of such data (see “Analysis of epegRNA phenotypes”).

#### Cloning of epegRNA libraries

##### Self-targeting library (lDS004)

A two-step cloning process was used. First, the Twist oligo pool was PCR amplified using Phusion Plus polymerase (ThermoFisher), 0.5 μM forward primer (5′-GTATCCCTTGGAGAACCACCT), 0.5 μM reverse primer (5′-CAGACGTGTGCTCTTCCGAT), and 0.1 pmol resuspended oligo pool with the following conditions: 1 cycle of 1 min at 98°C; 15 cycles of 15 s at 98°C, followed by 15 s at 60°C, followed by 45 s at 72°C; 1 cycle of 10 min at 72°C; 10°C hold. PCR products were purified using Machery-Nagel NucleoSpin Gel and PCR Clean-up kit per manufacturer protocol and quantified via Nanodrop. Vector backbone pAC025 was subjected to a BstXI-BamHI double restriction digest, followed by column cleanup. NEB Hifi DNA assembly was used to assemble the amplified library pool and digested vector in a 1:3 vector:insert ratio at 50°C for 1 hr. After SPRI purification, assembled products were transformed into electrocompetent cells (Endura) using a MicroPulser (BioRad). SOC media was added (for a total of 1.2 mL) and the transformation mixture was incubated at 37°C for 1 hr. The cells were then grown for 14 hr at 37°C in a 500 mL culture with LB broth and 100 μg mL^-1^ carbenicillin, and plasmids were extracted from the resulting cultures. To assess intermediate library coverage and quality, epegRNA cassettes and target regions were amplified for validation sequencing using flanking 5′ primer (5′-AATGATACGGCGACCACCGAGATCTACACGCACAAAAGGAAACTCACCCT) and 3′ indexing primer (5′-CAAGCAGAAGACGGCATACGAGATNNNNNNNNGTGACTGGAGTTCAGACGTGTGCTCTTC) with the following program: 1 cycle of 30 s at 98°C; 10 cycles of 10 s at 98°C, followed by 20 s at 65°C, followed by 20 s at 72°C; 1 cycle of 2 min at 72°C; 10°C hold. Sequencing was performed on Illumina MiSeq at 500X coverage (see “Sequencing”). Notably, sequencing revealed that epegRNA identities and their accompanying target regions with barcodes became uncoupled in ∼15% of reads, which we hypothesize may be due to the substantial homologous portions within and between each oligo. These uncoupled epegRNA-target site pairs were filtered from downstream analysis (see “Analysis of prime editing efficiencies”).

To complete the cloning, the intermediate library was digested with Esp3I enzyme (NEB R0734S) at 37°C for 6 hr and gel purified. The epegRNA scaffold sequence (5′-GTTTAAGAGCTATGCTGGAAACAGCATAGCAAGTTTAAATAAGGCTAGTCCGTTATCAACTTGAAAAAGTGGCACCGAGTCGGTGC)^60^ was synthesized with flanking reversed Esp3I sites (5′-CGTCTCGGTTT and 5′-GTGCTGAGACG) as a gene fragment by IDT and amplified by PCR using Phusion polymerase, 0.5 μM forward primer (5′-TCACAACTACACCAGAAGCCAC), 0.5 μM reverse primer (5′-GCTGGCAACACTTTGACGAAGA), and 0.1 pmole resuspended gene fragment with the following program: 1 cycle of 30 s at 98°C; 25 cycles of 10 s at 98°C, followed by 10 s at 58°C, followed by 15 s at 72°C; 1 cycle of 5 min at 72°C; 10°C hold. The amplified scaffold was purified by column cleanup and digested with Esp3I at 37°C for 6 hr. After column cleanup, the purified scaffold insert (2 ng) was ligated with the digested initial plasmid library vector (200 ng) using T4 ligase at 16°C overnight. After SPRI purification, ligated products were transformed into Endura electrocompetent cells as above. Final library quality was assessed via sequencing as above, with 90% of library elements occurring within a 6.1X range and a Gini coefficient of 0.26 (Figure S2A).

##### StopPR (lAC002)

As with the construction of lDS004, we used a two-step cloning process. First, the Twist oligo pool was PCR amplified using Phusion HSII HF (ThermoFisher), 0.4 μM forward primer (5′-CACCAGAAGCCACCTTGTTG), 0.4 μM reverse primer (5′-CTGTGTTGGTCTCCCGCG), and 10 ng resuspended oligo pool with the following program: 1 cycle of 30 s at 98°C; 6 cycles of 10 s at 98°C, followed by 20 s at 65°C, followed by 10 s at 72°C; 1 cycle of 5 min at 72°C; 10°C hold. Products from multiple PCR reactions were aggregated and purified using SPRI. Vector backbone pAC026 was subjected to a BstXI-BlpI (NEB R0585S) double digest at 37°C for 4 hr followed by SPRI purification, BsmBI-v2 (NEB R0739S) digest at 55°C for 6 hr, and final SPRI purification. Amplified oligo pool was double digested with BstXI and BsaI-v2 (NEB R3733S) restriction enzymes at 37°C for 4 hr and purified through column clean-up. Digested oligo pool and vector backbone were ligated using T4 DNA Ligase (NEB) at room temperature for 45 min and purified using SPRI. Transformation using electrocompetent Endura cells proceeded as described above, and library quality was assessed via sequencing. epegRNA cassettes were amplified for validation sequencing using primers as above for lDS004. Sequencing was performed on Illumina NovaSeq at 600X coverage (see “Sequencing”).

To complete the cloning, the intermediate library was digested with BsmBI-v2 enzyme at 55°C for 4 hr and SPRI purified. PCR amplification and purification of the epegRNA scaffold proceeded as above. Purified PCR product was digested with BsmBI-v2 at 55°C overnight, followed by SPRI purification. The purified scaffold insert (2 ng) was ligated with the digested intermediate plasmid library vector (200 ng) using T4 DNA Ligase (NEB) at room temperature for 45 min. After SPRI purification, ligated products were transformed into Endura electrocompetent cells and final library quality was assessed via sequencing as above. StopPR exhibited moderate skew resulting from missing elements (Gini coefficient of 0.35, with 90% of analyzed library elements present within a 57X range). After filtering lowly represented epegRNAs (see “Analysis of stop codon phenotypes”), we retained 84% of originally designed epegRNAs with well-distributed representation (Gini coefficient of 0.26, 90% of analyzed library elements present within a 5X range).

#### Production of lentivirus

Lentivirus production was performed for each library using the same process. HEK293T cells (14E6) were seeded in a 150-mm cell culture dish with DMEM. Plasmids pALD-Rev-A (1 μg, Aldevron), pALD-GagPol-A (1 μg, Aldevron), pALD-VSV-G-A (2 μg, Aldevron), and the transfer vector (15 μg) were mixed with Opti-MEM I Reduced Serum Medium (Gibco) and TransIT-LT1 (Mirus MIR 2300) transfection reagent, and co-transfected into cells. At 12-14 hr post-transfection, 1X ViralBoost reagent (ALSTEM) was added to cells, and at 48 hr post-transfection, lentivirus-containing supernatant was collected and stored at −80°C. To determine viral titer, serial dilutions of virus (500-0 μL) were transduced into K562 cells with 8 mg mL^-1^ polybrene. Titer was calculated 48 hr post-transduction based on the percent BFP fluorescent cells.

#### Arrayed endogenous site editing

The lentiviral pegRNAs and epegRNAs (*tevopreQ_1_*) targeting HEK3 and DNMT1 endogenous sites were transduced separately, each into a total of 0.6E6 cells for PE2, PEmax, and PEmaxKO stable cell lines in triplicate, at an MOI of 0.7. Cells were spun at 1000 x g for 2 hr in the presence of 8 mg mL^-1^ polybrene before incubating in a humidified incubator. Puromycin was added 72 hr post-transduction to deplete untransduced cells. To maintain coverage, cells were kept at a minimum of 2.5E7 cells per replicate, at a density of 0.5-1.0E6 cells mL^-1^ (splitting as necessary). Editing lasted for 28 days post-transduction, with time point samples collected at days 7, 14, 21, and 28. Genomic DNA (gDNA) was extracted from harvested K562 cells by first treating with lysis buffer (10 μM Tris-HCl, pH 7.5; 0.05% SDS; 25μg/mL Proteinase K), then by incubating at 37°C for 90 min followed by heat inactivation at 80°C for 30 min.

Endogenous sites were amplified from gDNA using a two-step PCR. First, flanking 5′ and 3′ primers were used to amplify HEK3 and DNMT1 genomic sites. HEK3 was amplified with flanking 5′ primer (5′-CGCCCATGCAATTAGTCTATTTCTGC) and 3′ primer (5′-CTCTGGGTGCCCTGAGATCTTTT), with the following program: 1 cycle of 2 min at 98°C; 32 cycles of 10 s at 98°C, followed by 20 s at 69°C, followed by 30 s at 72°C; 1 cycle of 2 min at 72°C; 10°C hold. DNMT1 was amplified with flanking 5′ primer (5′-CACAACAGCTTCATGTCAGCCAAG) and 3′ primer (3′-CGTTTGAGGAGTGTTCAGTCTC), with the following program: 1 cycle of 2 min at 98°C; 32 cycles of 10 s at 98°C, followed by 20 s at 66°C, followed by 30 s at 72°C; 1 cycle of 2 min at 72°C; 10°C hold. Resulting PCR1 products were SPRI purified using 1.0X reactions. Then, 5′ (5′-AATGATACGGCGACCACCGAGATCTACACNNNNNNNNACACTCTTTCCCTACACGAC) and 3′ (5′- CAAGCAGAAGACGGCATACGAGATNNNNNNNNGTGACTGGAGTTCAGACGTGTGCTCTTC) indexing primers were used to amplify purified PCR1 products, with the following program: 1 cycle of 2 min at 98°C; 8 cycles of 10 s at 98°C, followed by 20 s at 65°C, followed by 30 s at 72°C; 1 cycle of 2 min at 72°C; 10°C hold. Sequencing was performed on Illumina MiSeq at 50,000X coverage (see “Sequencing”).

#### Pooled screening

##### Self-targeting library (lDS004)

The lentiviral library was transduced into a total of 5E7 cells for both PEmax and PEmaxKO stable cell lines in replicate, at an MOI of 0.7 to achieve >10,000X coverage of the number of oligonucleotides. Cells were spun at 1000 x g for 2 hr in the presence of 8 mg mL^-1^ polybrene before incubating in a humidified multitron. 1 μg mL^-1^ Puromycin was added 72 hr post-transduction to deplete untransduced cells. To maintain coverage, cells were kept at a minimum of 2.5E7 cells per replicate (>10,000X coverage), at a density of 0.5-1.0E6 cells mL^-1^ (splitting as necessary). Screening lasted for 28 days post-transduction, with time point samples (12,500-25,000X representation) collected at days 7, 14, 21, and 28. gDNA was extracted from harvested K562 cells using the NucleoSpin Blood XL kit (Macherey Nagel). Subsequently, gDNA was treated with RNase A and purified by ethanol precipitation. epegRNA-target cassettes were PCR amplified using 5′ flanking primer (5′-AATGATACGGCGACCACCGAGATCTACACGCACAAAAGGAAACTCACCCT) and 3′ indexing primer (5′-CAAGCAGAAGACGGCATACGAGATNNNNNNNNGTGACTGGAGTTCAGACGTGTGCTCTTC).

Each 100 μL reaction contained 10 μg of genomic DNA, 1 μM primers, and 50 μL of NEBNext Ultra II Q5 Master Mix, and was run with the following program: 1 cycle of 1 min at 98°C; 22 cycles of 10 s at 98°C, followed by 30 s at 67°C, followed by 45 s at 72°C; 1 cycle of 5 min at 72°C; 10°C hold. Resulting PCR products from each sample were pooled and SPRI purified using 0.85-0.56X double-sided reactions.

##### StopPR (lAC002)

The lentiviral library was transduced into a total of 4.1E8 cells for PEmaxKO stable cell line in replicate, at an MOI of 0.7 to achieve >500X coverage of the number of oligonucleotides. Cells were spun at 1000xg for 2 hr in the presence of 8 mg mL^-1^ polybrene before incubating in a humidified multitron. 1 μg mL^-1^ Puromycin was added 72 hr post-transduction to deplete untransduced cells. To maintain coverage, cells were kept at a minimum of 4.5E8 cells per replicate (>1,500X coverage), at a density of 0.5-1.0E6 cells mL^-1^ (splitting as necessary). Screening lasted for 28 days post-transduction, with time point samples (1,250-2,000X representation) collected at days 7, 14, and 28. gDNA extraction and PCR amplification of epegRNA cassettes proceeded as above, under the following conditions: 1 cycle of 30s at 98°C; 22 cycles of 10s at 98°C, followed by 20s at 65°C, followed by 20s at 72°C; 1 cycle of 2 min at 72°C; 10°C hold. Resulting PCR products from each sample were pooled and SPRI purified using 0.85-0.56X double-sided reactions.

#### Sequencing

##### Endogenous sites

Sequencing was performed on an Illumina MiSeq with 10% phiX spike-in with single reads: I1 = 8nt, i7 index read; I2 = 8nt, i5 index read; R1 = 300nt, endogenous sequence. Standard Illumina primers were used for all reads.

##### Self-targeting library (lDS004)

Sequencing was performed on an Illumina MiSeq with 5% phiX spike-in with paired-end reads: I1 = 6nt, i7 index read; I2 = 0nt, i5 index read; R1 = 144nt, epegRNA spacer and extension; R2 = 68nt, target sequence and barcode. Custom primers were used for R1 (5′-GTGTGTTTTGAGACTATAAGTATCCCTTGGAGAACCACCTTGTTG), and standard Illumina primers were used for remaining reads.

##### StopPR (lAC002)

Sequencing was performed on an Illumina NovaSeq with 25% phiX spike-in with paired-end reads: I1 = 8nt, i7 index read; I2 = 0nt, i5 index read; R1 = 28nt, epegRNA spacer; R2 = 102nt, epegRNA extension. Custom primers were used for R1 as in sequencing of lDS004, and standard Illumina primers were used for remaining reads.

### Statistical Analysis

#### Analysis of prime editing efficiencies

##### Endogenous sites

To analyze sequencing data, we first used CRISPRessoBatch^63^ to align reads to HEK3 and DNMT1 reference endogenous sequences (inputted as -- amplicon_seq) based on spacer sequences (inputted as --guide_seq). Both min_average_read_quality and min_bp_quality_or_N arguments were set to 30, otherwise default parameters were used. The CRISPRessoBatch quantification window was positioned to include 20 nt on both sides of the Cas9(H840A) nick site (40 nt total window size). Custom Python scripts were used to further process aligned reads from CRISPRessoBatch (contained in allele frequency tables): first, to account for the presence of known SNPs at the endogenous targets in K562 cells, we allowed either A/G at the position 11 nt upstream of the nick site and either A/G at the position 9 nt downstream of the nick site for the HEK3 reference, and for the DNMT1 reference, we allowed either A/G at the position 3 nt upstream of the nick site. Second, we also considered nts assigned to “N” by CRISPRessoBatch, which likely arise due to sequencing errors, as reference (no edit or errors). We then collapsed reads into alignment bins accordingly. Reads were classified as either precise edit (only variant was the intended edit), no edit (same as reference sequence), or error (contained a variant that was not the intended edit), and reported efficiencies describe percentage of: (number of reads with the classified edit)/(number of reads that align to the amplicon).

##### Self-targeting library (lDS004)

Our self-targeting library was analyzed using a custom three-stage pipeline (Figure S2B):

In the first stage, each read was assigned to an epegRNA identity (unique to each epegRNA-target pair) by aligning components of the epegRNA (contained on Read 1) and target (contained on Read 2) to reference indices (*i.e.*, spacer through the end of epegRNA extension for Read 1, target sequence through the barcode for Read 2) using bwa mem.^64^ Read pairs with low mapping quality (≤ 5) or with recombination between the two reads were removed, and remaining reads were assigned to groups based on their epegRNA identities to enable parallel processing.

In stage two, the 45 nt target sites for each epegRNA-target pair were extracted, collapsed, and analyzed to determine observed editing outcomes. First, we extracted the part of the read that matched the reference target site with at least 60% of bases. As we have a 45 nt target site, outcomes with 18 or more nucleotide differences from the reference would have been discarded (defining an upper limit on observed indel lengths). Next, barcodes were extracted from reads by identifying the portion of reads that matched the expected barcode with no more than 8 mismatches, then any reads with errors in the barcodes (3 or more mismatched bases) were filtered to ensure that target sites matched epegRNA identities. Then, reads were collapsed to “outcomes” by identifying all reads with the same sequence. Outcomes that occurred at very low frequencies (0.1% or 10 total reads, whichever was higher) were filtered. We reasoned that the latter set of outcomes likely represented PCR or sequencing errors rather than edits introduced by prime editing. To deal with other outcomes likely containing systematic errors from low sequencing quality, we developed and applied the following algorithm: for each outcome, the mean sequencing quality score was calculated at each base; if the average quality was below 15 and the base did not match the reference sequence, it was corrected. This process was used sparingly, correcting a median of 33 reads per epegRNA-target pair across all four time points. After base correction, outcomes were globally aligned to their reference target sites, and variants (substitutions, insertions, and deletions) were called for each outcome. Each outcome was associated with zero (reference, no edits made) or more variants and classified as no edit (same as reference), precise edit (only variant is the intended edit), or error (contains a variant that isn’t the intended edit).

In stage three, all outcomes associated with individual epegRNA identities across all time points were aggregated into one file, and the resulting individual files were concatenated for analysis. Any pairs with fewer than 50 reads at any of the four collected timepoints were removed from analysis, with a unified set of epegRNA-target pairs analyzed for both cell lines.

#### Analysis of epegRNA phenotypes

To analyze deep sequencing data from the StopPR library, we used custom Python scripts to exactly match sequencing reads to epegRNA spacer and extension sequences. Excluded from reported statistics throughout the paper were pairs of spacer/codon-matched stop and synonymous epegRNAs for which either epegRNA converted a stop codon to a different stop codon, targeted a nonessential gene, or erroneously specified an edit in a noncoding region (found after updating validation code). These constituted a small minority of epegRNA pairs (580 total). Notably, this set of excluded epegRNAs included 68 epegRNAs (designed as synonymous) targeting the intronic base directly adjacent to 3′ exon boundaries; this small number of epegRNAs with unintended targets was used in the section, “Unbiased identification of splice site variants”. Additionally, we filtered any pairs of spacer/codon-matched stop and synonymous epegRNAs for which either epegRNA had fewer than 200 reads at day 7 (23,604). At day 14 and day 28, a pseudocount of 10 was added to all read counts to account for epegRNAs that had fully dropped out of the population. Enrichment of each epegRNA both at *Z* = day 14 and *Z* = day 28 was calculated as follows, where *Z _0_*= day 7:

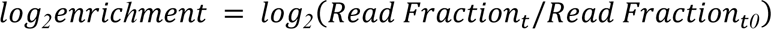

Enrichment was then normalized by subtracting the median enrichment of negative control epegRNAs (NC, non-targeting controls), resulting in our final growth phenotype measurement:

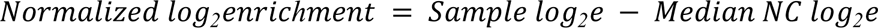

Phenotypes per epegRNA were averaged across replicates for both day 14 and 28, and all epegRNA phenotypes were converted to Z-scores by dividing them by the standard deviation of the non-targeting control epegRNA phenotypes. A phenotype induction cutoff was set as two standard deviations below the mean enrichment of non-targeting controls (*i.e.*, a score of Z < −2) based on previous literature.^7^ To determine a per-gene (or “gene-level”) stop epegRNA growth phenotype, the top two epegRNAs with the absolute largest stop epegRNA phenotypes for each gene were averaged.

#### Multiple linear regression model

To investigate the effects of different epegRNA design choices on phenotypic outcomes, we restricted our analysis to all stop epegRNAs which targeted a codon where phenotype induction was observed by at least one epegRNA. Subsetting the data in this manner isolated edits for which we had reasonable evidence that edit installation could induce phenotype. We reasoned that, in these cases, features other than the edit itself would determine differences in phenotype induction. This set of 51,279 stop epegRNAs was used to create a multiple linear regression model with the following features to predict day 14 phenotypes: edit distance from cut site (1-20 bp), edit length (1 bp, 2 bp, or 3 bp), edit installed (174 possibilities as no epegRNA specifying a CCT>TAA edit induced a phenotype), starting codon (59 possibilities), stop codon installed (TAG, TGA, TAA), PBS (11 nt, 13 nt) and RTT length (10 nt, 12 nt, 15 nt, 20 nt), spacer orientation relative to gene (sense or antisense), edit location within gene body (0-100%), and edit located within last exon of transcript (yes or no). Discrete features (starting codon, stop codon installed, substitution type, spacer orientation, last exon) were given numerical encodings through the use of 10-fold target encoding which, together with the coefficients from the resulting model, enabled a ranking of the relative importance of each category within the different features. We opted to use a target encoding approach to keep the dimensionality of our model low, as it directly replaces categorical features with their phenotypic mean. RTT length and edit position were given additional quadratic terms in the model to adjust from the observed preference of 15 nt RTT length and edits within the PAM region (Figures 4A and S4A). After encoding, all features were scaled to Z-scores by subtracting the mean and dividing by the standard deviation of each feature, and then the model was fit (Table S4).

#### ePRIDICT evaluation

We used ePRIDICT^57^ to generate chromatin favorability scores for prime editing for each stop epegRNA that survived filters in our StopPR library. For a small number of edits (639), ePRIDICT was missing needed chromatin features and thus did not generate scores, leaving a set of 101,857 stop epegRNAs targeting 15,008 codons for analysis. We defined a codon-level ePRIDICT score as the average ePRIDICT score from all targeted genomic positions within the same codon, and subsequently defined codons with score > 50 as having a favorable chromatin context, and those with score < 35 as having an unfavorable chromatin context, following thresholds for “high” and “low” scores defined in the original publication.^57^

#### Statistical testing and reproducibility

To compare top two stop epegRNA Z-scores between binnings of K562 CRISPRi phenotype, substitution position type, substitution length, gene body insertion location, and stop codon installed, we used a one-way analysis of variance (ANOVA) followed by two-sided Tukey’s post hoc test. To compare top two stop epegRNA enrichment values between binarized features including RTT and PBS lengths, spacer orientation relative to gene, and installation in the last exon, we used a two-sample t-test. When comparing all sense and antisense stop epegRNAs targeting the same substitution, we used a two-sample t-test. Effect size analysis was performed using the cohen.d function from the effsize R package with default parameters.^65^ For all analyses, NS p ≥ 0.05, * p < 0.05, ** p < 0.01, *** p < 0.001.

## References

1. Bick, A.G., Metcalf, G.A., Mayo, K.R., Lichtenstein, L., Rura, S., Carroll, R.J., Musick, A., Linder, J.E., Jordan, I.K., Nagar, S.D., et al. (2024). Genomic data in the All of Us Research Program. Nature, 1–7. 10.1038/s41586-023-06957-x.

2. Stenson, P.D., Mort, M., Ball, E.V., Chapman, M., Evans, K., Azevedo, L., Hayden, M., Heywood, S., Millar, D.S., Phillips, A.D., et al. (2020). The Human Gene Mutation Database (HGMD®): optimizing its use in a clinical diagnostic or research setting. Hum. Genet. 139, 1197–1207. 10.1007/s00439-020-02199-3.

3. Sahni, N., Yi, S., Taipale, M., Fuxman Bass, J.I., Coulombe-Huntington, J., Yang, F., Peng, J., Weile, J., Karras, G.I., Wang, Y., et al. (2015). Widespread Macromolecular Interaction Perturbations in Human Genetic Disorders. Cell 161, 647–660. 10.1016/j.cell.2015.04.013.

4. Stratton, M.R., Campbell, P.J., and Futreal, P.A. (2009). The cancer genome. Nature 458, 719–724. 10.1038/nature07943.

5. Schmidt, R., Ward, C.C., Dajani, R., Armour-Garb, Z., Ota, M., Allain, V., Hernandez, R., Layeghi, M., Xing, G., Goudy, L., et al. (2023). Base-editing mutagenesis maps alleles to tune human T cell functions. Nature, 1–8. 10.1038/s41586-023-06835-6.

6. Cuella-Martin, R., Hayward, S.B., Fan, X., Chen, X., Huang, J.-W., Taglialatela, A., Leuzzi, G., Zhao, J., Rabadan, R., Lu, C., et al. (2021). Functional interrogation of DNA damage response variants with base editing screens. Cell 184, 1081–1097.e19. 10.1016/j.cell.2021.01.041.

7. Hanna, R.E., Hegde, M., Fagre, C.R., DeWeirdt, P.C., Sangree, A.K., Szegletes, Z., Griffith, A., Feeley, M.N., Sanson, K.R., Baidi, Y., et al. (2021). Massively parallel assessment of human variants with base editor screens. Cell 184, 1064-1080.e20. 10.1016/j.cell.2021.01.012.

8. Cornu, T.I., and Cathomen, T. (2007). Targeted Genome Modifications Using Integrase-deficient Lentiviral Vectors. Mol. Ther. 15, 2107–2113. 10.1038/sj.mt.6300345.

9. Woods, N.T., Baskin, R., Golubeva, V., Jhuraney, A., De-Gregoriis, G., Vaclova, T., Goldgar, D.E., Couch, F.J., Carvalho, M.A., Iversen, E.S., et al. (2016). Functional assays provide a robust tool for the clinical annotation of genetic variants of uncertain significance. NPJ Genomic Med. 1, 16001. 10.1038/npjgenmed.2016.1.

10. Giacomelli, A.O., Yang, X., Lintner, R.E., McFarland, J.M., Duby, M., Kim, J., Howard, T.P., Takeda, D.Y., Ly, S.H., Kim, E., et al. (2018). Mutational processes shape the landscape of TP53 mutations in human cancer. Nat. Genet. 50, 1381–1387. 10.1038/s41588-018-0204-y.

11. Kotler, E., Shani, O., Goldfeld, G., Lotan-Pompan, M., Tarcic, O., Gershoni, A., Hopf, T.A., Marks, D.S., Oren, M., and Segal, E. (2018). A Systematic p53 Mutation Library Links Differential Functional Impact to Cancer Mutation Pattern and Evolutionary Conservation. Mol. Cell 71, 178–190.e8. 10.1016/j.molcel.2018.06.012.

12. Findlay, G.M., Boyle, E.A., Hause, R.J., Klein, J.C., and Shendure, J. (2014). Saturation editing of genomic regions by multiplex homology-directed repair. Nature 513, 120–123. 10.1038/nature13695.

13. Findlay, G.M., Daza, R.M., Martin, B., Zhang, M.D., Leith, A.P., Gasperini, M., Janizek, J.D., Huang, X., Starita, L.M., and Shendure, J. (2018). Accurate classification of BRCA1 variants with saturation genome editing. Nature 562, 217–222. 10.1038/s41586-018-0461-z.

14. Koblan, L.W., Arbab, M., Shen, M.W., Hussmann, J.A., Anzalone, A.V., Doman, J.L., Newby, G.A., Yang, D., Mok, B., Replogle, J.M., et al. (2021). Efficient C•G-to-G•C base editors developed using CRISPRi screens, target-library analysis, and machine learning. Nat. Biotechnol. 39, 1414–1425. 10.1038/s41587-021-00938-z.

15. Chen, L., Hong, M., Luan, C., Gao, H., Ru, G., Guo, X., Zhang, D., Zhang, S., Li, C., Wu, J., et al. (2023). Adenine transversion editors enable precise, efficient A•T-to-C•G base editing in mammalian cells and embryos. Nat. Biotechnol., 1–13. 10.1038/s41587-023-01821-9.

16. Sánchez-Rivera, F.J., Diaz, B.J., Kastenhuber, E.R., Schmidt, H., Katti, A., Kennedy, M., Tem, V., Ho, Y.-J., Leibold, J., Paffenholz, S.V., et al. (2022). Base editing sensor libraries for high-throughput engineering and functional analysis of cancer-associated single nucleotide variants. Nat. Biotechnol. 40, 862–873. 10.1038/s41587-021-01172-3.

17. Chen, P.J., and Liu, D.R. (2023). Prime editing for precise and highly versatile genome manipulation. Nat. Rev. Genet. 24, 161–177. 10.1038/s41576-022-00541-1.

18. Komor, A.C., Kim, Y.B., Packer, M.S., Zuris, J.A., and Liu, D.R. (2016). Programmable editing of a target base in genomic DNA without double-stranded DNA cleavage. Nature 533, 420–424. 10.1038/nature17946.

19. Gaudelli, N.M., Komor, A.C., Rees, H.A., Packer, M.S., Badran, A.H., Bryson, D.I., and Liu, D.R. (2017). Programmable base editing of A•T to G•C in genomic DNA without DNA cleavage. Nature 551, 464–471. 10.1038/nature24644.

20. Kurt, I.C., Zhou, R., Iyer, S., Garcia, S.P., Miller, B.R., Langner, L.M., Grünewald, J., and Joung, J.K. (2021). CRISPR C-to-G base editors for inducing targeted DNA transversions in human cells. Nat. Biotechnol. 39, 41–46. 10.1038/s41587-020-0609-x.

21. Tong, H., Wang, X., Liu, Y., Liu, N., Li, Y., Luo, J., Ma, Q., Wu, D., Li, J., Xu, C., et al. (2023). Programmable A-to-Y base editing by fusing an adenine base editor with an N-methylpurine DNA glycosylase. Nat. Biotechnol. 41, 1080–1084. 10.1038/s41587-022-01595-6.

22. Wang, J.Y., and Doudna, J.A. (2023). CRISPR technology: A decade of genome editing is only the beginning. Science 379, eadd8643. 10.1126/science.add8643.

23. Rees, H.A., and Liu, D.R. (2018). Base editing: precision chemistry on the genome and transcriptome of living cells. Nat. Rev. Genet. 19, 770–788. 10.1038/s41576-018-0059-1.

24. Anzalone, A.V., Koblan, L.W., and Liu, D.R. (2020). Genome editing with CRISPR-Cas nucleases, base editors, transposases and prime editors. Nat. Biotechnol. 38, 824–844. 10.1038/s41587-020-0561-9.

25. Anzalone, A.V., Randolph, P.B., Davis, J.R., Sousa, A.A., Koblan, L.W., Levy, J.M., Chen, P.J., Wilson, C., Newby, G.A., Raguram, A., et al. (2019). Search-and-replace genome editing without double-strand breaks or donor DNA. Nature 576, 149–157. 10.1038/s41586-019-1711-4.

26. Erwood, S., Bily, T.M.I., Lequyer, J., Yan, J., Gulati, N., Brewer, R.A., Zhou, L., Pelletier, L., Ivakine, E.A., and Cohn, R.D. (2022). Saturation variant interpretation using CRISPR prime editing. Nat. Biotechnol. 40, 885–895. 10.1038/s41587-021-01201-1.

27. Ren, X., Yang, H., Nierenberg, J.L., Sun, Y., Chen, J., Beaman, C., Pham, T., Nobuhara, M., Takagi, M.A., Narayan, V., et al. (2023). High-throughput PRIME-editing screens identify functional DNA variants in the human genome. Mol. Cell 83, 4633–4645.e9. 10.1016/j.molcel.2023.11.021.

28. Chardon, F.M., Suiter, C.C., Daza, R.M., Smith, N.T., Parrish, P., McDiarmid, T., Lalanne, J.-B., Martin, B., Calderon, D., Ellison, A., et al. (2023). A multiplex, prime editing framework for identifying drug resistance variants at scale. 10.1101/2023.07.27.550902.

29. Gould, S.I., Wuest, A.N., Dong, K., Johnson, G.A., Hsu, A., Narendra, V.K., Atwa, O., Levine, S.S., Liu, D.R., and Sánchez Rivera, F.J. (2024). High-throughput evaluation of genetic variants with prime editing sensor libraries. Nat. Biotechnol., 1–15. 10.1038/s41587-024-02172-9.

30. Kim, Y., Oh, H.-C., Lee, S., and Kim, H.H. (2023). Saturation resistance profiling of EGFR variants against tyrosine kinase inhibitors using prime editing. Preprint at bioRxiv, 10.1101/2023.12.03.569825 10.1101/2023.12.03.569825.

31. Martyn, G.E., Montgomery, M.T., Jones, H., Guo, K., Doughty, B.R., Linder, J., Chen, Z., Cochran, K., Lawrence, K.A., Munson, G., et al. (2023). Rewriting regulatory DNA to dissect and reprogram gene expression. Preprint at bioRxiv, 10.1101/2023.12.20.572268 10.1101/2023.12.20.572268.

32. Wang, T., Wei, J.J., Sabatini, D.M., and Lander, E.S. (2014). Genetic Screens in Human Cells Using the CRISPR-Cas9 System. Science 343, 80–84. 10.1126/science.1246981.

33. Shalem, O., Sanjana, N.E., Hartenian, E., Shi, X., Scott, D.A., Mikkelsen, T.S., Heckl, D., Ebert, B.L., Root, D.E., Doench, J.G., et al. (2014). Genome-Scale CRISPR-Cas9 Knockout Screening in Human Cells. Science 343, 84–87. 10.1126/science.1247005.

34. Konermann, S., Brigham, M.D., Trevino, A.E., Joung, J., Abudayyeh, O.O., Barcena, C., Hsu, P.D., Habib, N., Gootenberg, J.S., Nishimasu, H., et al. (2015). Genome-scale transcriptional activation by an engineered CRISPR-Cas9 complex. Nature 517, 583–588. 10.1038/nature14136.

35. Gilbert, L.A., Horlbeck, M.A., Adamson, B., Villalta, J.E., Chen, Y., Whitehead, E.H., Guimaraes, C., Panning, B., Ploegh, H.L., Bassik, M.C., et al. (2014). Genome-Scale CRISPR-Mediated Control of Gene Repression and Activation. Cell 159, 647–661. 10.1016/j.cell.2014.09.029.

36. Nuñez, J.K., Chen, J., Pommier, G.C., Cogan, J.Z., Replogle, J.M., Adriaens, C., Ramadoss, G.N., Shi, Q., Hung, K.L., Samelson, A.J., et al. (2021). Genome-wide programmable transcriptional memory by CRISPR-based epigenome editing. Cell 184, 2503–2519.e17. 10.1016/j.cell.2021.03.025.

37. Chen, P.J., Hussmann, J.A., Yan, J., Knipping, F., Ravisankar, P., Chen, P.-F., Chen, C., Nelson, J.W., Newby, G.A., Sahin, M., et al. (2021). Enhanced prime editing systems by manipulating cellular determinants of editing outcomes. Cell 184, 5635–5652.e29. 10.1016/j.cell.2021.09.018.

38. Yan, J., Oyler-Castrillo, P., Ravisankar, P., Ward, C.C, Levesque, S., Jing, Y., Simpson, D., Zhao, A., Li, H., Yan, W., et al. Improving prime editing with an endogenous small RNA-binding protein. Nature, in press. 10.1038/s41586-024-07259-6.

39. Ferreira da Silva, J., Oliveira, G.P., Arasa-Verge, E.A., Kagiou, C., Moretton, A., Timelthaler, G., Jiricny, J., and Loizou, J.I. (2022). Prime editing efficiency and fidelity are enhanced in the absence of mismatch repair. Nat. Commun. 13, 760. 10.1038/s41467-022-28442-1.

40. Nelson, J.W., Randolph, P.B., Shen, S.P., Everette, K.A., Chen, P.J., Anzalone, A.V., An, M., Newby, G.A., Chen, J.C., Hsu, A., et al. (2022). Engineered pegRNAs improve prime editing efficiency. Nat. Biotechnol. 40, 402–410. 10.1038/s41587-021-01039-7.

41. Choi, J., Chen, W., Suiter, C.C., Lee, C., Chardon, F.M., Yang, W., Leith, A., Daza, R.M., Martin, B., and Shendure, J. (2022). Precise genomic deletions using paired prime editing. Nat. Biotechnol. 40, 218–226. 10.1038/s41587-021-01025-z.

42. Li, X., Chen, W., Martin, B.K., Calderon, D., Lee, C., Choi, J., Chardon, F.M., McDiarmid, T., Kim, H., Lalanne, J.-B., et al. (2023). Chromatin context-dependent regulation and epigenetic manipulation of prime editing. BioRxiv Prepr. Serv. Biol., 2023.04.12.536587. 10.1101/2023.04.12.536587.

43. Lahue, R.S., Au, K.G., and Modrich, P. (1989). DNA Mismatch Correction in a Defined System. Science 245, 160–164.

44. Su, S.S., Lahue, R.S., Au, K.G., and Modrich, P. (1988). Mispair specificity of methyl-directed DNA mismatch correction in vitro. J. Biol. Chem. 263, 6829–6835. 10.1016/S0021-9258(18)68718-6.

45. Shen, M.W., Arbab, M., Hsu, J.Y., Worstell, D., Culbertson, S.J., Krabbe, O., Cassa, C.A., Liu, D.R., Gifford, D.K., and Sherwood, R.I. (2018). Predictable and precise template-free CRISPR editing of pathogenic variants. Nature 563, 646–651. 10.1038/s41586-018-0686-x.

46. Kim-Yip, R.P., McNulty, R., Joyce, B., Mollica, A., Chen, P.J., Ravisankar, P., Law, B.K., Liu, D.R., Toettcher, J.E., Ivakine, E.A., et al. (2024). Efficient prime editing in two-cell mouse embryos using PEmbryo. Nat. Biotechnol., 1–9. 10.1038/s41587-023-02106-x.

47. Allen, F., Crepaldi, L., Alsinet, C., Strong, A.J., Kleshchevnikov, V., De Angeli, P., Páleníková, P., Khodak, A., Kiselev, V., Kosicki, M., et al. (2019). Predicting the mutations generated by repair of Cas9-induced double-strand breaks. Nat. Biotechnol. 37, 64–72. 10.1038/nbt.4317.

48. Kim, H.K., Kim, Y., Lee, S., Min, S., Bae, J.Y., Choi, J.W., Park, J., Jung, D., Yoon, S., and Kim, H.H. (2019). SpCas9 activity prediction by DeepSpCas9, a deep learning–based model with high generalization performance. Sci. Adv. 5, eaax9249. 10.1126/sciadv.aax9249.

49. Arbab, M., Shen, M.W., Mok, B., Wilson, C., Matuszek, Ż., Cassa, C.A., and Liu, D.R. (2020). Determinants of Base Editing Outcomes from Target Library Analysis and Machine Learning. Cell 182, 463–480.e30. 10.1016/j.cell.2020.05.037.

50. Kim, H.K., Yu, G., Park, J., Min, S., Lee, S., Yoon, S., and Kim, H.H. (2021). Predicting the efficiency of prime editing guide RNAs in human cells. Nat. Biotechnol. 39, 198–206. 10.1038/s41587-020-0677-y.

51. Mathis, N., Allam, A., Kissling, L., Marquart, K.F., Schmidheini, L., Solari, C., Balázs, Z., Krauthammer, M., and Schwank, G. (2023). Predicting prime editing efficiency and product purity by deep learning. Nat. Biotechnol. 41, 1151–1159. 10.1038/s41587-022-01613-7.

52. Yu, G., Kim, H.K., Park, J., Kwak, H., Cheong, Y., Kim, D., Kim, J., Kim, J., and Kim, H.H. (2023). Prediction of efficiencies for diverse prime editing systems in multiple cell types. Cell 186, 2256–2272.e23. 10.1016/j.cell.2023.03.034.

53. 53. DepMap, B. (2022). DepMap 22Q2 Public. (figshare). 10.6084/M9.FIGSHARE.19700056.V2 10.6084/M9.FIGSHARE.19700056.V2.

54. Horlbeck, M.A., Gilbert, L.A., Villalta, J.E., Adamson, B., Pak, R.A., Chen, Y., Fields, A.P., Park, C.Y., Corn, J.E., Kampmann, M., et al. Compact and highly active next-generation libraries for CRISPR-mediated gene repression and activation. eLife 5, e19760. 10.7554/eLife.19760.

55. Cohen, J. (1992). Statistical Power Analysis. Curr. Dir. Psychol. Sci. 1, 98–101. 10.1111/1467-8721.ep10768783.

56. Lykke-Andersen, S., and Jensen, T.H. (2015). Nonsense-mediated mRNA decay: an intricate machinery that shapes transcriptomes. Nat. Rev. Mol. Cell Biol. 16, 665–677. 10.1038/nrm4063.

57. Mathis, N., Allam, A., Tálas, A., Benvenuto, E., Schep, R., Damodharan, T., Balázs, Z., Janjuha, S., Schmidheini, L., Böck, D., et al. (2023). Predicting prime editing efficiency across diverse edit types and chromatin contexts with machine learning (Molecular Biology) 10.1101/2023.10.09.561414.

58. Sibley, C.R., Blazquez, L., and Ule, J. (2016). Lessons from non-canonical splicing. Nat. Rev. Genet. 17, 407–421. 10.1038/nrg.2016.46.

59. Zhang, S., Samocha, K.E., Rivas, M.A., Karczewski, K.J., Daly, E., Schmandt, B., Neale, B.M., MacArthur, D.G., and Daly, M.J. (2018). Base-specific mutational intolerance near splice sites clarifies the role of nonessential splice nucleotides. Genome Res. 28, 968–974. 10.1101/gr.231902.117.

60. Chen, B., Gilbert, L.A., Cimini, B.A., Schnitzbauer, J., Zhang, W., Li, G.-W., Park, J., Blackburn, E.H., Weissman, J.S., Qi, L.S., et al. (2013). Dynamic imaging of genomic loci in living human cells by an optimized CRISPR/Cas system. Cell 155, 1479– 1491. 10.1016/j.cell.2013.12.001.

61. Doench, J.G., Fusi, N., Sullender, M., Hegde, M., Vaimberg, E.W., Donovan, K.F., Smith, I., Tothova, Z., Wilen, C., Orchard, R., et al. (2016). Optimized sgRNA design to maximize activity and minimize off-target effects of CRISPR-Cas9. Nat. Biotechnol. 34, 184–191. 10.1038/nbt.3437.

62. Cunningham, F., Allen, J.E., Allen, J., Alvarez-Jarreta, J., Amode, M.R., Armean, I.M., Austine-Orimoloye, O., Azov, A.G., Barnes, I., Bennett, R., et al. (2022). Ensembl 2022. Nucleic Acids Res. 50, D988–D995. 10.1093/nar/gkab1049.

63. Clement, K., Rees, H., Canver, M.C., Gehrke, J.M., Farouni, R., Hsu, J.Y., Cole, M.A., Liu, D.R., Joung, J.K., Bauer, D.E., et al. (2019). CRISPResso2 provides accurate and rapid genome editing sequence analysis. Nat. Biotechnol. 37, 224–226. 10.1038/s41587-019-0032-3.

64. Li, H., and Durbin, R. (2009). Fast and accurate short read alignment with Burrows-Wheeler transform. Bioinforma. Oxf. Engl. 25, 1754–1760. 10.1093/bioinformatics/btp324.

65. 65. Torchiano, M. (2016). Effsize - a package for efficient effect size computation. (Zenodo). 10.5281/ZENODO.1480624 10.5281/ZENODO.1480624.

