## Supplementary Information for "A benchmarked, high-efficiency prime editing platform for multiplexed dropout screening"

### Supplementary Figures

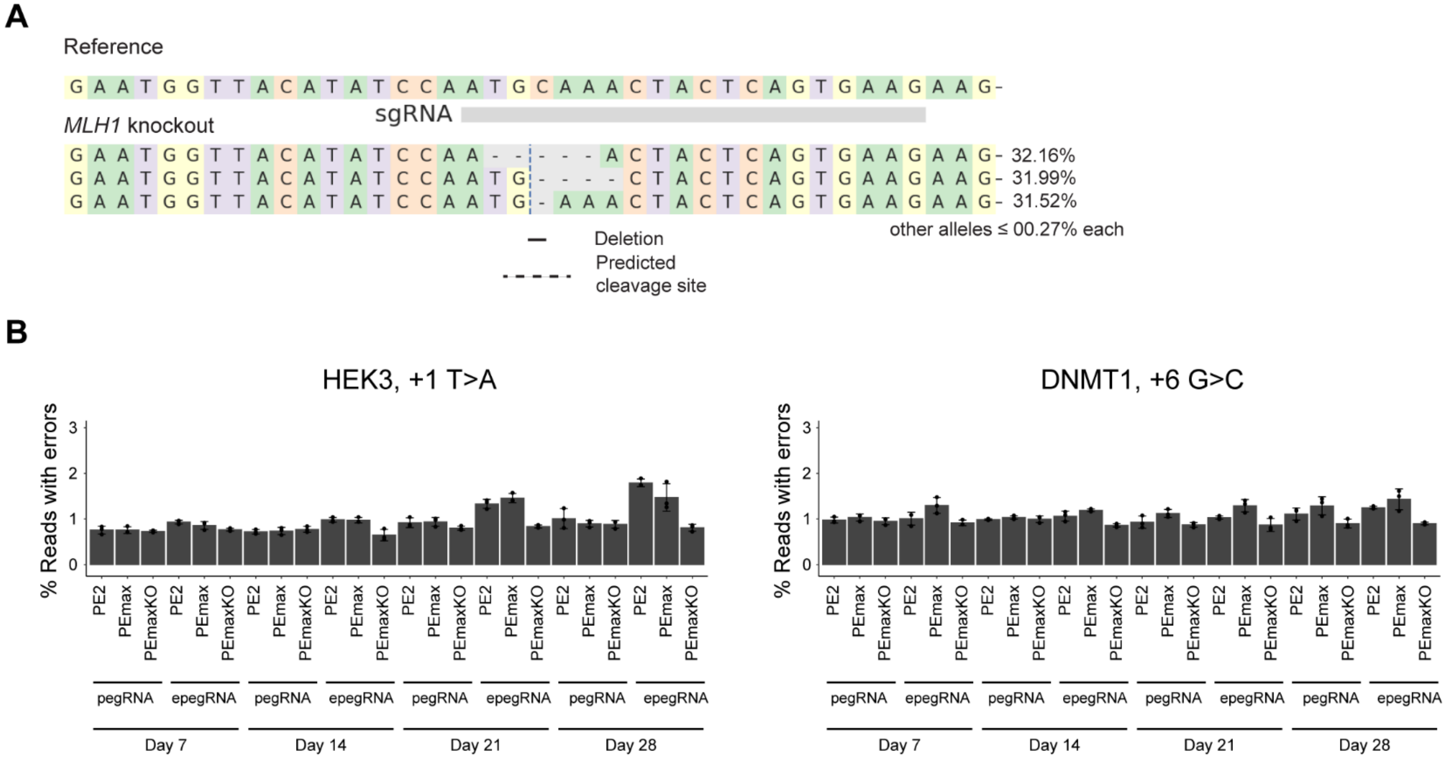


**Figure S1. Continuous prime editing at two endogenous loci produces very low frequencies of on-target, unwanted editing (errors), related to Figure 1.**

(A) Sequences and frequencies of alleles observed at the targeted *MLH1* locus in our PEmaxKO cell line.

(B) Percentages of sequencing reads with unintended, on-target edits (errors) from indicated conditions. Corresponding frequencies of precise editing presented in Figure 1C. Data and error bars represent mean +/- s.d. (n=3 independent biological replicates).

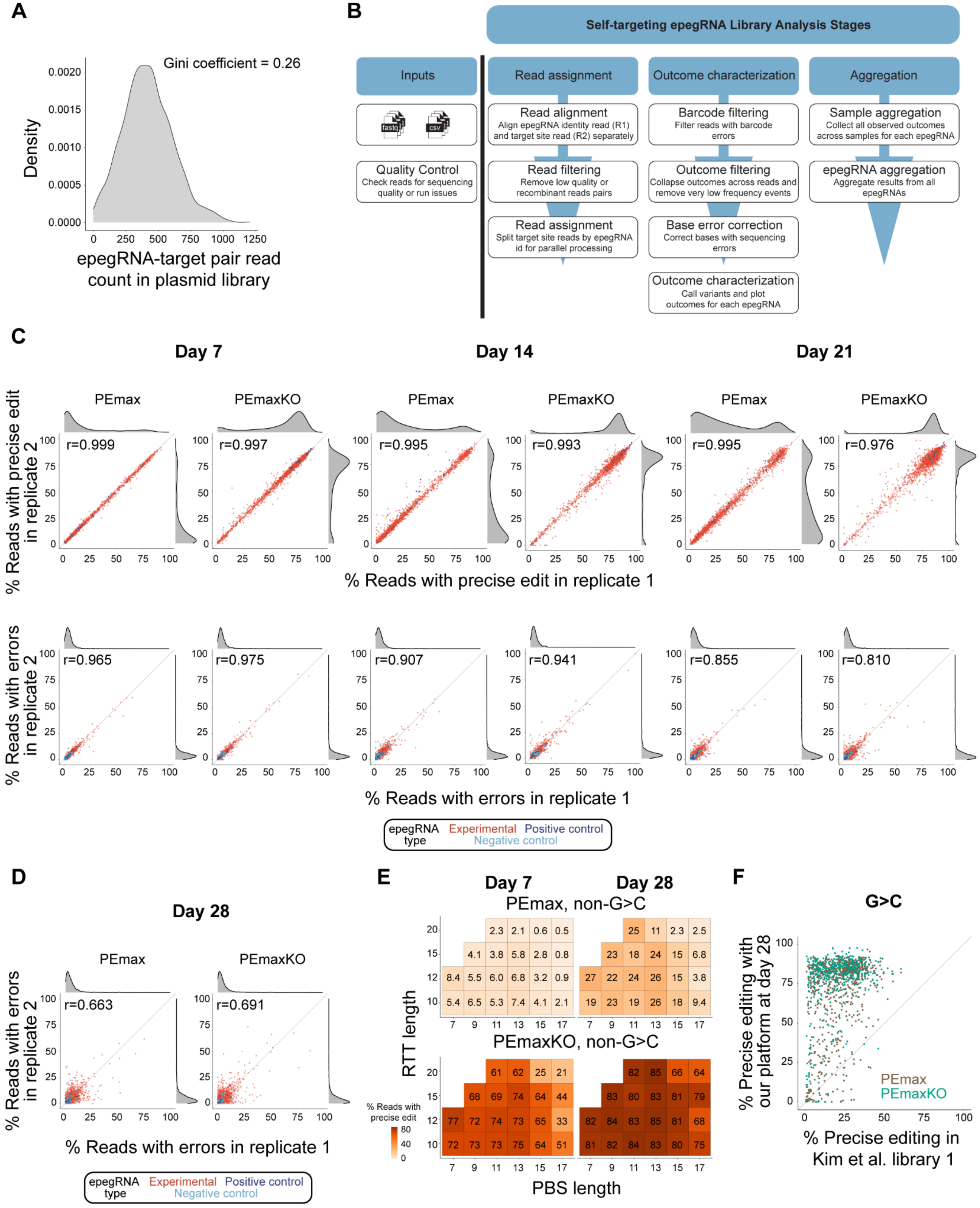
A
Gini coefficient = 0.26
Density
epegRNA-target pair read count in plasmid library
B
Self-targeting epegRNA Library Analysis Stages
Inputs
fastq
csv
Quality Control
Check reads for sequencing quality or run issues
Read assignment
Read alignment
Align epegRNA identity read (R1) and target site read (R2) separately
Read filtering
Remove low quality or recombinant reads pairs
Read assignment
Split target site reads by epegRNA id for parallel processing
Outcome characterization
Barcode filtering
Filter reads with barcode errors
Outcome filtering
Collapse outcomes across reads and remove very low frequency events
Base error correction
Correct bases with sequencing errors
Outcome characterization
Call variants and plot outcomes for each epegRNA
Aggregation
Sample aggregation
Collect all observed outcomes across samples for each epegRNA
epegRNA aggregation
Aggregate results from all epegRNAs
C
Day 7
Day 14
Day 21
PEmax
PEmaxKO
r=0.999
r=0.997
r=0.995
r=0.993
r=0.995
r=0.976
% Reads with precise edit in replicate 2
% Reads with precise edit in replicate 1
r=0.965
r=0.975
r=0.907
r=0.941
r=0.855
r=0.810
% Reads with errors in replicate 2
% Reads with errors in replicate 1
epegRNA type
Experimental
Positive control
Negative control
D
Day 28
PEmax
PEmaxKO
r=0.663
r=0.691
% Reads with errors in replicate 2
% Reads with errors in replicate 1
epegRNA type
Experimental
Positive control
Negative control
E
Day 7
Day 28
PEmax, non-G>C
PEmaxKO, non-G>C
RTT length
PBS length
% Reads with precise edit
F
G>C
% Precise editing with our platform at day 28
% Precise editing in Kim et al. library 1
PEmax
PEmaxKO

**Figure S2. Analysis of data from self-targeting sensor screens demonstrates that precise editing with low error rates is reproducible over one month of continuous prime editing, related to Figure 2.**

(A) Count distribution of epegRNA-target reads from self-targeting sensor library (plasmid). Gini coefficient indicates the degree of inequality on a scale from 0 (perfectly equal representation among all elements) to 1 (only one element represented). Typically, libraries with coefficients less than 0.30 are reasonably well distributed. Here, 90% of library elements occurred within a 6.1X range.

(B) Computational pipeline used to analyze results from self-targeting sensor screen. Analysis happened in three stages, as described in the Methods. Briefly, sequencing reads were assigned to an individual epegRNA-target pair, then editing outcomes were characterized for each pair as unedited, precise, or errors, and finally results were aggregated.

(C) Percentages of sequencing reads from sensor targets containing the precise edit (top) and errors (below) from two replicates of screens performed as indicated. Dots represent data from individual epegRNA-target pairs. Correlation between replicates (Pearson's r) indicated. Density plots on top and side show data distribution for replicate 1 and 2, respectively.

(D) As in C, but only for errors from indicated screens, collected on day 28.

(E) Heatmap depicting median, replicated-averaged percentages of sequencing reads from sensor targets containing the precise edit (only for non-G>C edits) for different RTT and PBS lengths of experimental epegRNAs (percentages indicated). Data from cells collected on indicated days from indicated screens, shown for RTT/PBS combinations that were used to target at least five sensor targets.

(F) Comparison between G>C editing efficiencies achieved with our platform (*y*-axis, replicate-averaged) and a previous report (*x*-axis) that used transiently expressed PE2 and pegRNAs in HEK293T cells over five days of editing.[50] Dots indicate edits evaluated in both studies, with efficiencies resulting from our PEmax and PEmaxKO screens shown in brown and green, respectively.

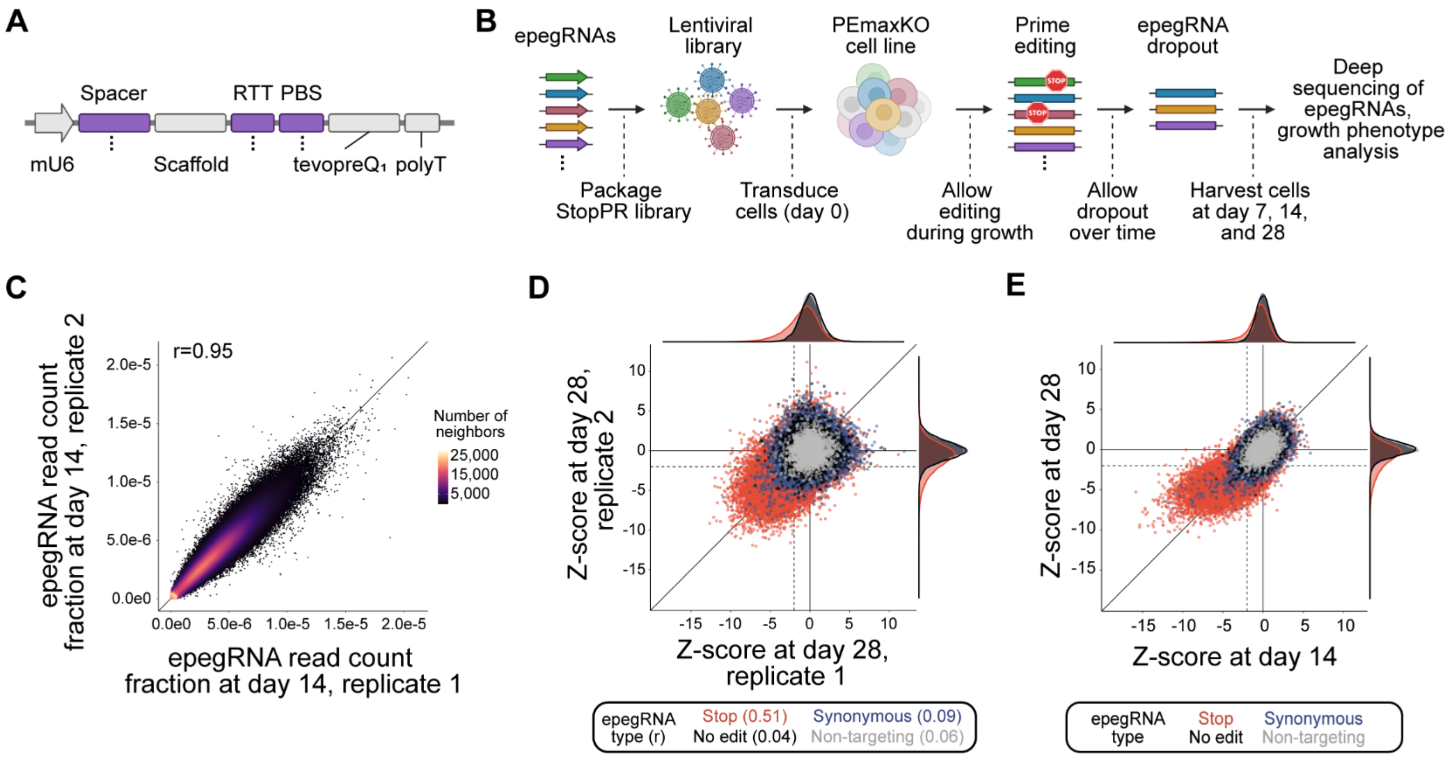


**Figure S3. Growth phenotypes from stop epegRNAs accumulate but become noisier over time, related to Figure 3.**

(A) Schematic of epegRNA expression cassette used for StopPR screen. Regions indicated with purple varied coordinately across the library (as denoted by dots). mU6, modified mouse U6 promoter; RTT, reverse transcriptase template; PBS, primer binding site.

(B) Schematic of workflow for StopPR screen. Briefly, epegRNAs were transduced into PEmaxKO cells and cell populations were grown for 28 days and sampled intermittently to evaluate prime editing-induced growth phenotypes.

(C) Normalized read counts (relative to total sequencing reads) for each epegRNA from independent biological replicates of StopPR screen sampled at day 14. Data points colored by density, indicated by number of neighbors. Correlation between replicates (Pearson's r) indicated.

(D) Growth phenotypes for epegRNAs from independent biological replicates of StopPR screen collected 28 days post-transduction. Dotted lines denote phenotype cutoffs ($Z < -2$). Correlation (Pearson's r) indicated for each epegRNA type.

(E) Growth phenotypes (averaged across replicates) from StopPR screen collected 14 and 28 days post-transduction. Dotted lines denote phenotype cutoffs ($Z < -2$).

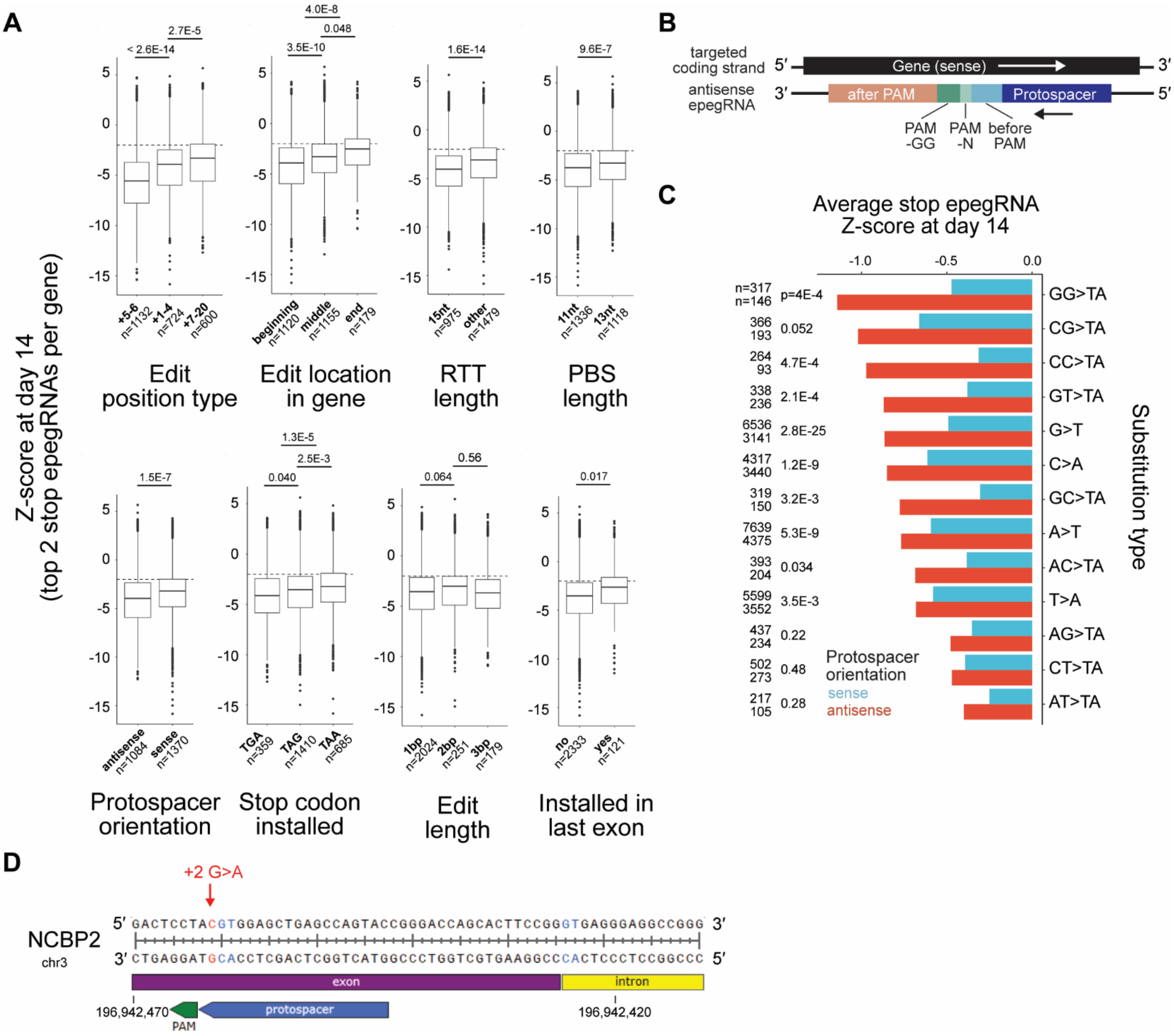

A
Z-score at day 14
(top 2 stop epegRNAs per gene)
< 2.6E-14
2.7E-5
+5-6
n=1132
+1-4
n=724
+7-20
n=600
Edit
position type
3.5E-10
4.0E-8
0.048
beginning
n=1120
middle
n=1155
end
n=179
Edit location
in gene
1.6E-14
15nt
n=975
other
n=1479
RTT
length
9.6E-7
11nt
n=1336
13nt
n=1118
PBS
length
1.5E-7
antisense
n=1084
sense
n=1370
Protospacer
orientation
0.040
1.3E-5
2.5E-3
TGA
n=359
TAG
n=1410
TAA
n=685
Stop codon
installed
0.064
0.56
1bp
n=2024
2bp
n=251
3bp
n=179
Edit
length
0.017
no
n=2333
yes
n=121
Installed in
last exon
B
targeted
coding strand
5′
Gene (sense)
3′
antisense
epegRNA
3′
after PAM
Protospacer
5′
PAM
-GG
PAM
-N
before
PAM
C
Average stop epegRNA
Z-score at day 14
-1.0
-0.5
0.0
n=317
n=146
p=4E-4
GG>TA
366
193
0.052
CG>TA
264
93
4.7E-4
CC>TA
338
236
2.1E-4
GT>TA
6536
3141
2.8E-25
G>T
4317
3440
1.2E-9
C>A
319
150
3.2E-3
GC>TA
7639
4375
5.3E-9
A>T
393
204
0.034
AC>TA
5599
3552
3.5E-3
T>A
437
234
0.22
AG>TA
502
273
0.48
CT>TA
217
105
0.28
AT>TA
Protospacer
orientation
sense
antisense
Substitution type
D
+2 G>A
NCBP2
chr3
5′ GACTCCTACGTGGAGCTGAGCCAGTACCGGGACCAGCACTTCCGGGTGAGGGAGGCCGGG 3′
3′ CTGAGGATGCACCTCGACTCGGTCATGGCCCTGGTCGTGAAGGCCCACTCCCTCCGGCCC 5′
exon
intron
196,942,470
PAM
protospacer
196,942,420

**Figure S4. epegRNA and endogenous target features vary in their impact on phenotype, related to Figure 4.**

(A) Growth phenotypes for stop epegRNAs (top two per gene) from StopPR screen sampled from day 14, binned by indicated features. Feature categories and analyzed epegRNAs comprise those used for effect size analysis in Figure 4A. For features with two categories, a two-sample t-test was performed. For features with three categories, an ANOVA followed by Tukey post-hoc was performed. Resulting p-values indicated. Median and interquartile range (IQR) of the full set of epegRNAs used in this analysis are indicated. Whiskers extend 1.5*IQR past the upper and lower quartiles.

(B) Schematic of antisense protospacer orientation relative to gene. Colors as in Figure 4B.

(C) Average growth phenotypes for stop epegRNAs specifying individual substitution types targeted in our library with both sense (blue) and antisense (red) protospacers. Data from cells sampled from day 14 of StopPR screen. Numbers of stop epegRNAs and p-values from two-sample t-test significance denoted (left).

(D) Endogenous sequence in *NCBP2* targeted by three of the 69 synonymous epegRNAs with unexpectedly strong growth phenotypes ($Z < -5$ at day 14), with phenotype possibly due to cryptic splice site activation. Red indicates specified edit. Blue indicates splice donor motifs. Numbers beneath each track indicate genomic coordinates. Locus is oriented to show the coding sequence of *NCBP2* from 5′ to 3′, but as the gene is positioned on the negative strand, genomic coordinates increase in the opposite direction (right to left).

#### Supplementary Tables

**Table S1. Self-targeting sensor library design and results, related to Figures 2 and S2.** Information about the design of each epegRNA and its associated target site used in the self-targeting sensor screens, along with resulting editing and error rates at each timepoint. Specifically, Freq.EditOnly (proportion of reads with only the intended edit) was used when reporting precise editing rates, and Freq.TotalErrors (proportion of reads with at least one unintended edit) was used when reporting error rates. Additionally, Freq.TotalWildType (proportion of reads that were unedited) is included. Average columns report the indicated frequencies averaged across replicates for the same epegRNA-target pair for the relevant screen and sampled timepoint.

**Table S2. StopPR library design and results for control epegRNAs, related to Figures 3, S3, 4, and S4.** Information about the design of each control epegRNA (non-targeting, no edit, and synonymous) used in the StopPR screen, along with resulting growth phenotypes (reported as Z-scores) at each timepoint. Average columns report the indicated Z-score (either at day 14 or day 28) averaged across replicates for the same epegRNA.

**Table S3. StopPR library design and results for stop epegRNAs, related to Figures 3, S3, 4, and S4.** Information about the design of each stop epegRNA used in the StopPR screen, along with resulting growth phenotypes (reported as Z-scores) at each timepoint. Average columns report the indicated Z-score (either at day 14 or day 28) averaged across replicates for the same epegRNA.

**Table S4. Multilinear model for impact of epegRNA and endogenous target features, related to "Multiple linear regression model" in Methods.** Model coefficients and significance for the multilinear model discussed in the Results. Model encodings used for categorical variables are also provided.

| Publication | Cell line | Editor protein | Editing system | Stable editor | Stable editor delivery | epegRNA | MMR-deficient | Largest library size | Readout | Primary screening application |
|---|---|---|---|---|---|---|---|---|---|---|
| Erwood et al.[26] | HEK-293T (haploid-ized) | PE2 | PE3 | No | - | No | Yes (HEK-293T) | 117 | Endogenous editing | SGE of *NPC1* and *BRCA2* regions |
| Ren et al.[27] | MCF7 | PE2 | PE3 | Yes | Lentiviral transduction | Yes | No | 6,744 | epegRNA abundance | Dropout screen on *MYC* enhancer, clinical variants |
| Chardon et al.[28] | PC-9 | PEmax | PE2 | Yes | Piggybac | Yes | Yes (*MLH1* KO) | 3,825 | epegRNA abundance | Enrichment screen for TKI resistance of *EGFR* variants |
| Gould et al.[29] | A549 | PEmax | PE2 | Yes | Lentiviral transduction | Yes | No | 28,219 | Sensor editing | Dropout and enrichment screen on *TP53* variants |
| Kim and Oh et al.[30] | PC-9 | PEmax | PE2 | Yes | Lentiviral transduction | Yes | Yes (*MLH1* KO) | 1,845 | Endogenous editing | SGE of *EGFR* region, with TKI context |
| Martyn and Montgomery et al.[31] | THP-1 | PE2 | PE2 | Yes | Lentiviral transduction | Yes | No | 185 | Endogenous editing | Flow-FISH screen on *PPIF* promoter and enhancer |
| *This manuscript* | K562 | PEmax | PE2 | Yes | *AAVS1* knockin | Yes | Yes (*MLH1* KO) | 240,000 | epegRNA abundance | Dropout screen on pan-essential genes |

**Table S5. Comparison of features used in different phenotype-based prime editing screens.** Compiled information from different published or preprinted prime editing screen manuscripts. System components listed represent the most optimized or relevant combinations used in each publication, in respect to this work (additional cell lines, editing systems, *etc*. have been explored in some publications). PE3 refers to prime editing with an additional nicking guide RNA (ngRNA). Under Readout, sensor editing refers to self-targeting style screens which require additional sequencing of exogenous target sites, while endogenous editing refers to requiring additional sequencing of the endogenous locus. KO, knockout; SGE, saturation genome editing; TKI, tyrosine kinase inhibitors.
